# Insights into the *Klebsiella pneumoniae* adaptive response mechanisms to colistin exposure using a label-free quantitative proteomics approach

**DOI:** 10.64898/2026.03.26.714365

**Authors:** Sambit K. Dwibedy, Indira Padhy, Sushil K. Pathak, Saswat S. Mohapatra

## Abstract

The rise of MDR *Klebsiella pneumoniae* and its resistance to the last-resort antibiotic colistin poses a significant threat to global healthcare. While genomic studies have identified several resistance mutations, the transient proteomic shifts that occur during the initial exposure of sensitive strains to lethal antibiotic doses remain poorly characterised. In this study, we employed a label-free quantitative proteomics approach to investigate the protein expression profile of *K. pneumoniae* strain ATCC 13883 treated with colistin at its MIC. Membrane proteins were extracted at critical growth stages, and differentially abundant proteins (DAPs) were analysed using Gene Ontology and KEGG pathway enrichment analysis. Our proteomic analysis identified 718 DAPs (339 upregulated and 379 downregulated). The cellular response was characterised primarily by outer membrane remodelling and a significant upregulation of the capsule-associated kinase Wzc and the ArnBCADTEF operon, which facilitates lipid A modification with L-Ara4N moiety. Paradoxically, while RND-family efflux pumps (AcrAB) were significantly induced, the global activator RamA and major porins (OmpA, OmpX, LamB) were downregulated, possibly to minimise antibiotic entry. KEGG pathway enrichment analysis further revealed a synchronised metabolic shift, characterised by an intensified TCA cycle flux to fuel high-energy resistance processes despite a general slowdown in carbohydrate metabolism. Our findings demonstrate that *K. pneumoniae* responds to colistin stress through a rapid, multifaceted proteomic reorganisation involving charge neutralisation, structural reinforcement of the cell envelope, and metabolic re-routing. These results provide a molecular blueprint of the early adaptive response, identifying several proteins as potential therapeutic targets.

## Introduction

The emergence of multidrug-resistant (MDR) *Klebsiella pneumoniae* in hospital settings is a major healthcare concern in recent times [1]. Carbapenem-resistant *K. pneumoniae* (CRKP) strains are notorious for their association with multiple nosocomial infections, including bacteraemia, pneumonia, and urinary tract infection (UTI) in immunocompromised patients [2,3]. For the treatment of infections caused by the CRKP strains, polymyxins, a class of cationic antimicrobial peptides (CAMPs), are used [4,5]. Polymyxins, including the therapeutically effective polymyxin B and polymyxin E (colistin), are largely used against MDR Gram-negative bacteria (GNB) such as *Escherichia coli, K. pneumoniae, Pseudomonas aeruginosa*, and *Acinetobacter baumannii*, and are often considered as a last option against these formidable pathogens [4,5]. Polymyxins target the lipid A moiety of the lipopolysaccharide (LPS) present in the outer membrane (OM) of GNB and work by interfering with membrane permeability [5,6]. However, due to the extensive use of polymyxins over the last few decades, several strains of *K. pneumoniae* have developed resistance to it [1,7,8].

Resistance to polymyxins primarily arises from modification of the LPS in the bacterial OM. LPS modification in *K. pneumoniae* is mostly mediated by the two-component systems (TCS) such as PhoPQ, PmrAB, and CrrAB, which activate the *arnBCADTEF* operon, and the *eptA* gene encoding the phosphoethanolamine transferase enzyme [6,9]. The remodelling of lipid A involves two main modifications: the addition of 4-amino-4-deoxy-L-arabinose (L-Ara4N) and the addition of phosphoethanolamine (pEtN) [10]. The addition of L-Ara4N and pEtN moieties to lipid A attenuates its electrostatic interaction with the positively charged α, γ-diaminobutyric acid (Dab) residue of polymyxin, weakening the binding affinity of polymyxin, leading to the development of resistance [11]. Phosphoethanolamine transferases can also be encoded by mobile colistin resistance (*mcr*) genes carried on conjugative plasmids, conferring polymyxin resistance and further disseminating them [12]. While the mechanisms of bacterial resistance to polymyxin are largely understood, the global cellular response to polymyxin treatment remains incomplete.

Proteomics studies have the potential to provide a far more complete understanding of the cellular response to the antibiotic treatment and have been used successfully in recent times [13–17]. Proteomics investigations have often discovered novel proteins and pathways that can be targeted for future drug development. Specifically, in *K. pneumoniae*, studies have been performed to detect differentially expressed proteins in antibiotic-sensitive vs antibiotic-resistant strains [13,14,16–20]. However, most studies have used antibiotic-resistant strains and compared their proteomes with those of their sensitive counterparts. The study by Fleitas et al. (2024) used the laboratory evolution approach to develop a *K. pneumoniae* strain resistant to the antimicrobial peptide PaDBS1R1 and compared its proteome. Considering that sensitive and resistant strains are not genetically identical, they would certainly have distinct proteome signatures, as observed in such studies. However, limited data are available on proteomic changes in sensitive bacteria exposed to high antibiotic doses. Therefore, we argue that proteomic perturbations in the sensitive bacterial population exposed to lethal doses of the antibiotic would play a major role in the response mechanism and the subsequent development of the resistance phenotype, and these effects need to be studied.

In the present study, we sought to understand the global proteomic changes in the *K. pneumoniae* strain exposed to colistin (polymyxin E) by comparing differentially expressed proteins between treated and untreated conditions using a label-free quantitative proteomics approach. Our study has revealed that exposure to colistin leads to large-scale OM remodelling and the activation of the CAMP resistance pathway, which may contribute to the development of colistin resistance over time.

## Materials and methods

### Bacterial strains and reagents

The *K. pneumoniae* strain ATCC 13883 was used in this study [21]. The *E. coli* strain ATCC 25922 was included as a quality control strain in the antibiotic susceptibility assay. The strains were routinely grown in LB medium and Mueller–Hinton broth (MHB) as and when required. The culture media, reagents, antibiotic discs, and E-test strips were procured from HiMedia Laboratories, Mumbai, India. Stock solutions of colistin sulphate (PCT1142) (10 mg/ml) were prepared using Milli-Q water (Millipore). BugBuster HT Protein extraction reagent (Cat#70922) and Protease Inhibitor, EDTA-free tablet (Cat#A32955) were obtained from Merck and Thermo Scientific, respectively.

### Growth curve

The standard growth curve of *K. pneumoniae* was determined by growing the strain in 50 ml of LB broth at 37°C with shaking at 150 rpm. The optical density of the culture at 600 nm (OD_600_) was measured using the UV-vis spectrophotometer (Genesys 180, Thermo Scientific) every 1 hour for 12 hours. The experiment was performed in triplicate, and the mean and standard deviations were calculated. An OD_600_ vs. time curve was plotted to determine the different growth phases.

### Antibiotic susceptibility profile and the estimation of colistin MIC

The antibiotic sensitivity pattern of the *K. pneumoniae* strain was determined using the Kirby-Bauer disk diffusion test as described previously [21] following the Clinical and Laboratory Standards Institute (CLSI) guidelines [22].

The MIC of colistin against *K. pneumoniae* was determined by the E-test and broth microdilution (BMD) methods, as described previously [21], in accordance with CLSI guidelines [22]. Briefly, for the BMD assay, the bacterial strain was grown to an OD corresponding to a 0.5 McFarland standard, diluted 300-fold, and tested against a 2-fold serial dilution of colistin. Growth inhibition was assessed after 20 hours of incubation [21]. The MIC of colistin was also determined using the E-test method by placing the colistin E-strip on the MH agar plate previously spread with a 0.5 McFarland standard inoculum [21]. The result was interpreted after 20 hours of incubation by checking the intersection of the zone with the strip.

### Colistin treatment and time-kill assay

The impact of colistin on *K. pneumoniae* growth was assessed through an *in vitro* time-kill assay as described previously [15], with a few modifications. Initially, the strain was cultured in LB broth until the OD_600_ reached 0.4, corresponding to approximately 2.1 × 10^6^ CFU/ml. This OD value was chosen to represent a standardised point in the bacterial growth curve. The bacterial culture was divided into four experimental groups, each treated with colistin at 0 µg/ml, 1 µg/ml, 1.5 µg/ml, or 2 µg/ml, respectively. Samples were collected from each group at 0, 1, 2, and 4 hours to measure OD_600_ and viable cell count (CFU/ml). The number of viable bacteria was estimated by performing a 10-fold serial dilution of the sample and plating on LB agar plates in triplicate. A graph of log_10_(CFU/ml) value was plotted against time. The data obtained from the study were analysed using GraphPad Prism software (version 10.0.0 for Windows, GraphPad Software, Boston, Massachusetts, USA).

### Growth conditions and sample preparation for proteomics

*K. pneumoniae* was grown at 37°C with shaking at 150 rpm to an OD_600_ of 0.4 (mid-log phase). At this point, colistin at a final concentration of 2 µg/ml, corresponding to the MIC, was added, and growth continued. A parallel culture without colistin was also grown under the same conditions. Samples (35 ml) were withdrawn from the cultures at 1, 2, and 4 hours after antibiotic addition. Cells were harvested by centrifugation at 8000 g at 4°C for 15 min. The cells were stored at -20°C till further use.

### Cell lysis and extraction of the membrane proteins

The frozen bacterial pellets were thawed on ice, then treated with the cell lysis buffer (450 µl) composed of the BugBuster HT Protein extraction reagent (Merck, Millipore), mixed with one Protease Inhibitor, EDTA-free tablet (Thermo Scientific). The samples were mixed gently by tapping the tubes. The BugBuster-treated cell samples were incubated on a shaking platform at room temperature for 20 minutes. The lysates were transferred to microcentrifuge tubes and incubated for an additional 30 minutes at room temperature. The cell lysates were centrifuged at 15,500 g for 15 minutes at 4°C twice. The supernatant containing the total protein was carefully separated, transferred to a 15 ml centrifuge tube, and stored at -20°C.

The cell lysates were diluted to a final volume of 10 ml with TBS buffer containing 150 mM NaCl, 50 mM Tris-Cl (pH 7.6), and protease inhibitor. The samples were subjected to ultracentrifugation at 1,25,000 g, at 4°C for 45 minutes using a Beckman Coulter Optima XPM-100 ultracentrifuge. Post ultracentrifugation, the supernatant was discarded by inverting the centrifuge tubes. Any remaining supernatant was carefully removed using a micropipette. The pellet containing the total membrane protein fraction was resuspended in 300 µl of a buffer consisting of 150 mM NaCl, 50 mM EDTA (pH 7.6), and protease inhibitor. The pellets were completely resuspended in the buffer by vortexing. The samples were then stored at -20°C until further use.

The membrane protein concentration was estimated using a 96-well plate-based Bradford assay. Briefly, a standard curve was generated using serially diluted bovine serum albumin (BSA) with concentrations ranging from 0.625 to 10 µg/ml in TBS buffer. The diluted protein samples and BSA standards were loaded in triplicate into the 96-well microplate, followed by the addition of Bradford reagent (Bio-Rad) to each well. Plates were incubated at room temperature for 10 minutes. The absorbance was then measured at 595 nm using a 96-well plate reader (Bio-Rad). Protein concentrations of unknown samples were calculated by linear regression analysis from the BSA standard curve [23].

### Membrane protein profiling by SDS-PAGE

Equal amounts of protein (4 µg) from different samples were run on a 10% SDS-PAGE, and protein bands were detected by Coomassie Brilliant Blue staining. The membrane proteins were assessed for differential expression by visualising band intensities in the colistin-treated and untreated samples. Based on visual differences in band intensity, the protein bands were selected, and their molecular weights were determined using ImageJ software with reference to the molecular weight markers.

### Label-free quantitative proteomics analysis

The protein sample preparation, subsequent mass spectrometry, and analysis were performed at the Advanced Mass Spectrometry Facility of the Institute of Life Sciences, Bhubaneswar, using the proteomics platform’s methodologies, as described below.

### Sample preparation for proteomics studies

Six volumes of ice-cold LC-MS-grade acetone were added to the protein sample, and the mixture was incubated at -20°C overnight. The mixture was centrifuged at 8000 g for 5 minutes at 4°C. The supernatant was discarded carefully, and the resulting protein pellet was air-dried at room temperature to remove residual acetone. The pellet was then resuspended in a buffer containing 8 M urea and 50 mM ammonium bicarbonate.

The protein buffer exchange was conducted using an Amicon Ultra 0.5 ml, 3K centrifugal filter device (Millipore #UFC500324). The device was equilibrated with 200 µl of a 1 M urea-containing dissolution buffer, as per the manufacturer’s protocol. The assembled device was then centrifuged at 14,000 g for 20 minutes to prepare it for sample loading. The resulting flow-through was discarded. Subsequently, up to 500 µl of the sample was carefully added to the equilibrated device. The sample was mixed gently by inverting the tube. The sample was then centrifuged at 14,000 g for 30 minutes at least thrice, each time using fresh dissolution buffer, to achieve complete buffer exchange and the desired sample volume. After the final centrifugation step, the filter was inverted to collect the concentrated sample. The concentrated samples were stored at -20°C.

### Trypsin digestion

For the in-solution trypsin digestion, the protein samples were reconstituted in a solution containing 8 M urea and 50 mM ammonium bicarbonate at pH 7.5-8.0. A total of 100 µg of protein was prepared, and the final volume was adjusted to 100 µl using the same buffer. For the reduction of disulfide bonds, 5 µl of 200 mM dithiothreitol (DTT) was added to the mixture. The solution was then incubated for 45 minutes at room temperature in the dark. Subsequently, 20 µl of 200 mM iodoacetamide (IAA) was added to the mixture to alkylate cysteine residues, and the sample was incubated for an additional 45 minutes at room temperature in the dark. The urea concentration was then reduced to 0.5 M by adding a solution of 50 mM ammonium bicarbonate and 1 mM calcium chloride (pH 7.6). This dilution step was necessary to ensure optimal enzyme activity, as high urea concentrations can inhibit trypsin. Trypsin was added to the protein sample at a final ratio of 1:50 (trypsin to protein, w/w). The mixture was gently vortexed and incubated at 37°C overnight to facilitate complete digestion. To terminate the enzymatic reaction, formic acid was added until the pH was lowered to 3−4. The resulting peptides were then desalted using C18 spin columns for downstream analysis.

### LC-MS/MS analysis

To identify differentially expressed proteins, quantitative proteomics analysis by LC-MS/MS was performed on membrane protein samples from *K. pneumoniae* treated with colistin, along with untreated control samples. Protein samples were analysed in triplicate. Peptides were analysed using an Orbitrap Fusion Lumos Tribrid mass spectrometer (Thermo Scientific) coupled with a Vanquish Neo LC system operating in a “Trap and Elute” injection workflow. Samples were loaded at 100 µl/min and 800 bar before separation on a 50 cm analytical column with a 75 µm internal diameter. Chromatographic separation was achieved at a constant flow rate of 300 nl/min for 140 minutes, with a gradient from 5% to 95% phase B (80% acetonitrile and 0.1% formic acid).

MS was performed in positive ion mode using a static nano electrospray ionisation (NSI) source with a spray voltage of 1900 V and an ion transfer tube temperature of 275°C. MS^1^ spectra were acquired in the Orbitrap at a resolution of 60,000 across a scan range of m/z 350–2000. Data-dependent acquisition followed a “Top 20” format, where precursors with charge states 2–6 and an intensity above 5e^3^ were selected for HCD fragmentation using stepped normalised collision energies of 28, 30, and 31%. MS^2^ spectra were recorded in the Ion Trap at a rapid scan rate in centroid mode. Dynamic exclusion was applied for 20 seconds with a mass tolerance of 10 ppm to prevent redundant sampling of high-abundance precursors. The raw files generated in MS were analysed using Proteome Discoverer 3.1 (Thermo Scientific) with the search engine Sequest HT, using 0.01 FDR and a cut-off of 2 missed cleavages as input parameters, as described previously [24–26].

### Database search and analysis

The spectral data were searched against the reference proteomes for *K. pneumoniae* (UniProt ID: UP000000265; Taxon ID 272620) downloaded from UniProt (https://www.uniprot.org/) on January 22, 2025. Precursor and product ion tolerances were set at 10 ppm and 0.05 Da, respectively. Oxidation of methionine (+15.995 Da) was treated as a variable modification, and carbamidomethylation of cysteine (+57.021 Da) as a static modification. Peptide-spectrum matches (PSMs) were refined to achieve an FDR of 0.01. Protein quantification was performed by aggregating reporter ion counts from all corresponding PSMs. Statistically significant proteins were selected from the total identified proteins based on Student’s *t*-test scores. Based on the expression values in each sample, differentially abundant proteins (DAPs) were identified as having a fold change (colistin-treated/control) ≥1.5 for upregulation and ≤0.66 for downregulation. Furthermore, the fold change values of the identified DAPs were transformed to log_2_ fold change (log_2_FC). The statistically significant proteins were identified using an FDR-corrected *p*-value ≤ 0.05 [25,26].

The data were further processed using MetaboAnalyst 5.0 [27] to identify DAPs between control and colistin-treated samples. The normalised data were first subjected to principal component analysis (PCA) to visualise the overall sample distribution and identify potential outliers or clustering patterns between the control and antibiotic-treated groups. Results were visualised using volcano plots and heat maps (following hierarchical clustering) to illustrate expression patterns.

### Gene ontology (GO) and KEGG pathway enrichment analysis

Each protein in *K. pneumoniae* ATCC 13883 was annotated against the genome of the type strain *K. pneumoniae* MGH 78578 using homology. The gene ontology (GO) terms for MGH 78578 were retrieved from the UniProt database and assigned to the corresponding ATCC 13883 proteins. The GO and KEGG databases were used to annotate all identified proteins with functional and pathway information. Enrichment analyses for the GO and KEGG pathways were performed using the hypergeometric distribution method to identify biological functions and pathways enriched among the DAPs in the compared samples, using the web-based tools ShinyGo [28] and KEGG Mapper [29]. Entries with *p*-values < 0.05 were considered significant.

### Prediction of sub-cellular localisation

The subcellular localisation of the DAPs was predicted using a protein localisation predictor tool, PSORTb version 3.0.3 [30].

## Results

### Growth characteristics and antibiotic susceptibility profile

Previously, we performed an adaptive laboratory evolution (ALE) study using the *K. pneumoniae* strain ATCC 13883 and showed that the strain can develop resistance to colistin and polymyxin B within a very short time [21]. Moreover, the evolved strain exhibited stable resistance and heteroresistance phenotypes for more than 100 generations, even in the absence of antibiotic selection [21]. In this study, we explored proteomic changes in bacteria exposed to high doses of colistin to understand the global cellular response to antibiotic stress.

The *K. pneumoniae* strain exhibits a typical growth pattern in LB broth, with a log phase lasting approximately 3 hours (Fig. 1A). The strain was found to be susceptible to most antibiotics, including colistin and polymyxin B, whereas resistance was observed against four antibiotics, i.e., ampicillin, erythromycin, clindamycin, and rifampicin. The MIC of colistin, as determined by the E-test, was 1.5 μg/ml (Fig. 1B), whereas the BMD assay showed a MIC of 2 μg/ml (Fig. S1).

**Fig. 1.**
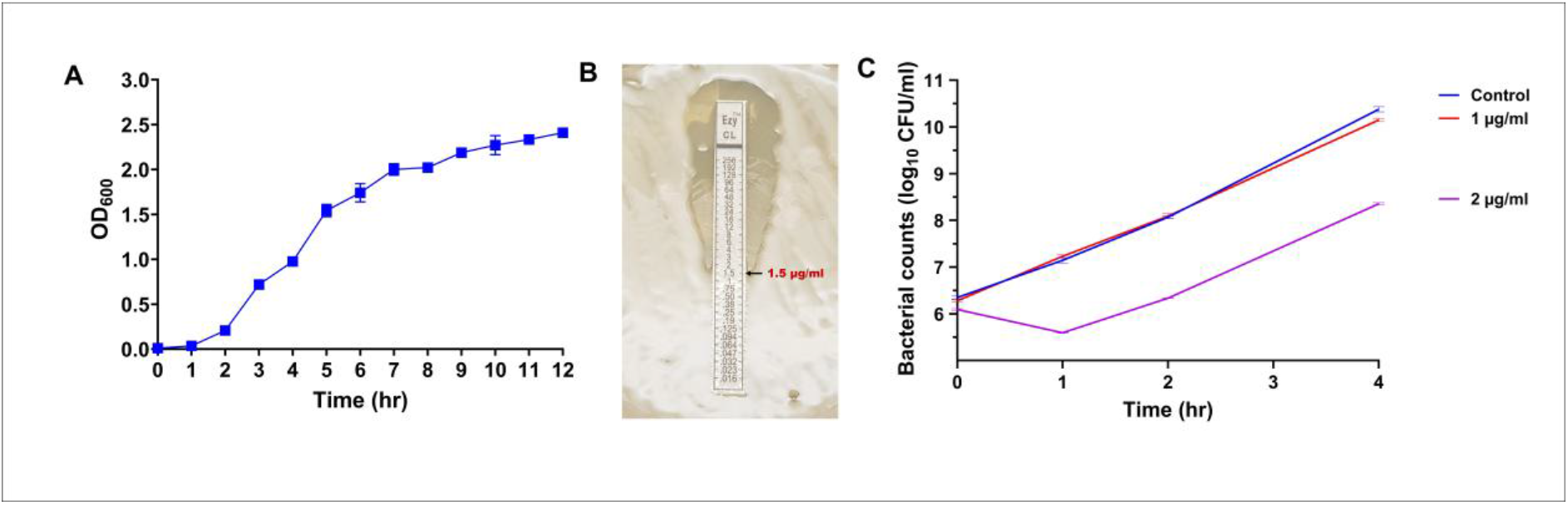
Growth characteristics and response to colistin treatment. (A) Growth kinetics of *K. pneumoniae* ATCC 13883 grown in LB broth for 12 hours, monitored by measuring the OD_600_. Error bars represent the standard deviation from triplicate experiments. (B) Determination of the colistin MIC using the E-test on Mueller-Hinton Agar (MHA). (C) Time-kill kinetics of *K. pneumoniae* exposed to colistin. The plot shows viable bacterial count (log_10_CFU/ml) over 4 hours following treatment with sub-MIC (1 μg/ml) and MIC (2 μg/ml) concentrations of colistin, compared with an untreated control.

### Impact of colistin treatment on *K. pneumoniae* growth

To understand the effect of colistin treatment on *K. pneumoniae* growth, we performed a time-kill assay using two different colistin concentrations-1 µg/ml (sub-MIC) and 2 µg/ml (MIC), as determined in the MIC assay. We tested the effect of the said antibiotic concentrations at different stages of bacterial growth (early log, mid-log, and late log). Colistin treatment at the early log phase (OD_600_ = 0.2) arrested the bacterial growth leading to population extinction, whereas its impact at the late log phase (OD_600_ = 0.7) was negligible, indicating an inoculum effect. However, treatment at the mid-log phase (OD_600_ = 0.4) showed a typical time-kill curve, where a substantial reduction in viable bacterial count was observed within the first hour (Fig. 1C), indicating a significant bactericidal effect of colistin. However, the trend reversed after the first hour, resulting in an increase in numbers at 2 hours and a significant increase in bacterial population by 4 hours. Considering the primary goal of the work to understand the proteome dynamics during the colistin treatment and the subsequent bacterial response to it, for subsequent experiments, we chose to treat the bacteria at mid-log phase with 2 µg/ml of colistin.

### Membrane proteome extraction and profiling

After the treatment of *K. pneumoniae* culture with colistin, total membrane proteins were extracted at three time points (1, 2, and 4 hr). Membrane proteins were also prepared from the culture grown without colistin treatment, which served as the control. Protein concentration was measured using the Bradford assay. The membrane proteins were profiled using the SDS-PAGE (Fig. S2). Membrane proteins from colistin-treated samples showed several differentially expressed protein bands compared with the control. Based on the distinct protein expression profile at 2 hours of colistin treatment, we selected this time point for subsequent quantitative proteomics.

### Label-free quantitative proteomics

This study employed quantitative proteomics using LC-MS/MS to investigate the key biological processes underlying the response to colistin treatment in *K. pneumoniae*. The PCA score plot showed that samples from each group clustered tightly within the 95% confidence interval, indicating that differences in protein profiles between the control and colistin-treated *K. pneumoniae* can be easily distinguished (Fig. 2A). Analysis of the membrane proteome of colistin-treated *K. pneumoniae* revealed that stress associated with high concentrations of antibiotics induces specific changes in the proteome. The data showed that colistin-treated and untreated *K. pneumoniae* exhibited distinct protein expression patterns, as visualised in a volcano plot (Fig. 2B). The thresholds for determining DAPs were set at a fold change (FC) ≥ 1.5 or ≤ 0.66 for up- and downregulated proteins, respectively, with a *p*-value ≤ 0.05. Out of the normalised 1071 proteins, 718 DAPs were found fitting into the above criteria (Suppl. data sheet 1), of which 379 were downregulated, and 339 were upregulated in the colistin-treated *K. pneumoniae* (Fig. 2B). Expression patterns of DAPs between samples are shown in the heatmap (Fig. 3). Details of the top DAPs are presented in Table 1.

**Table 1.**
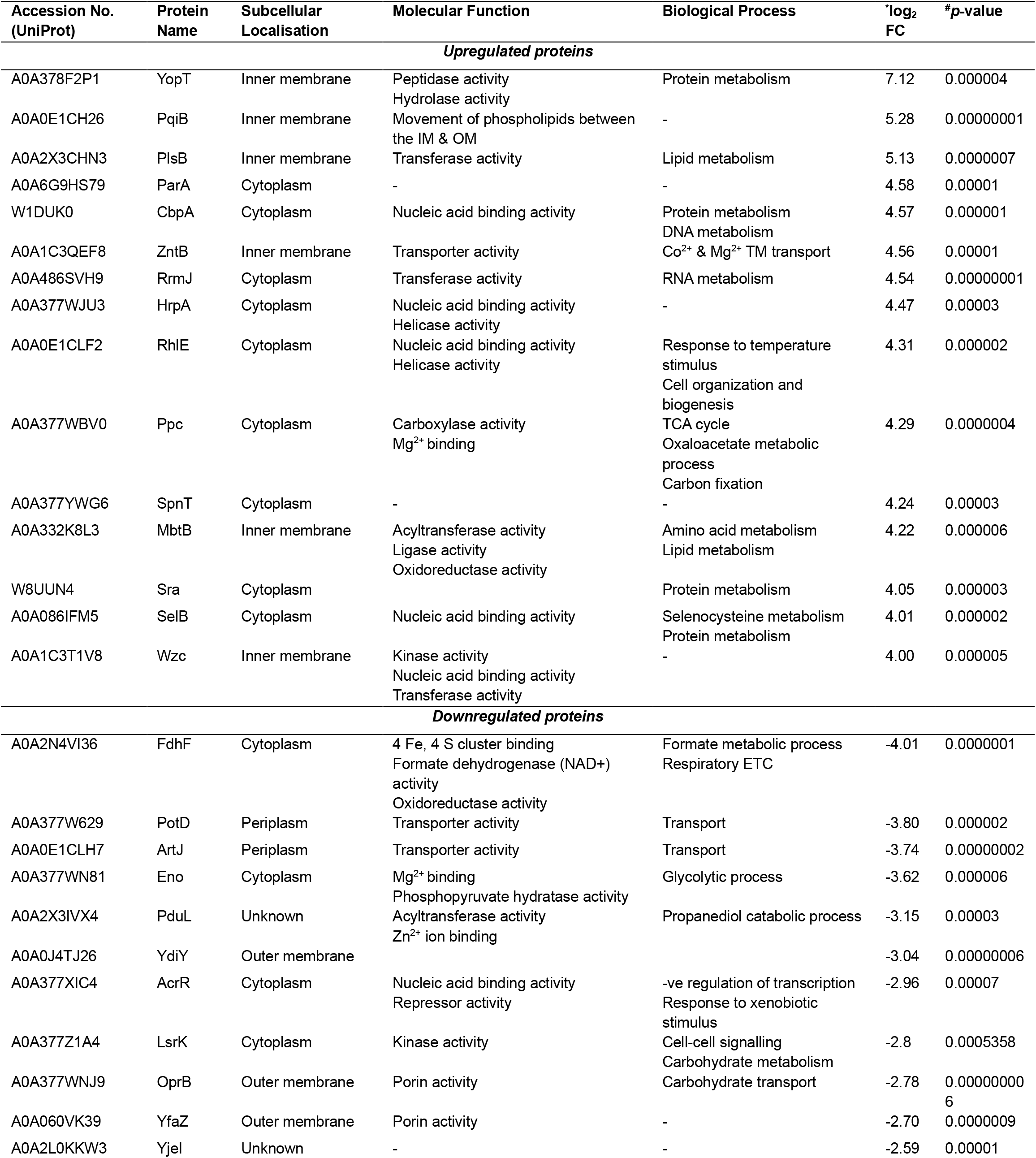
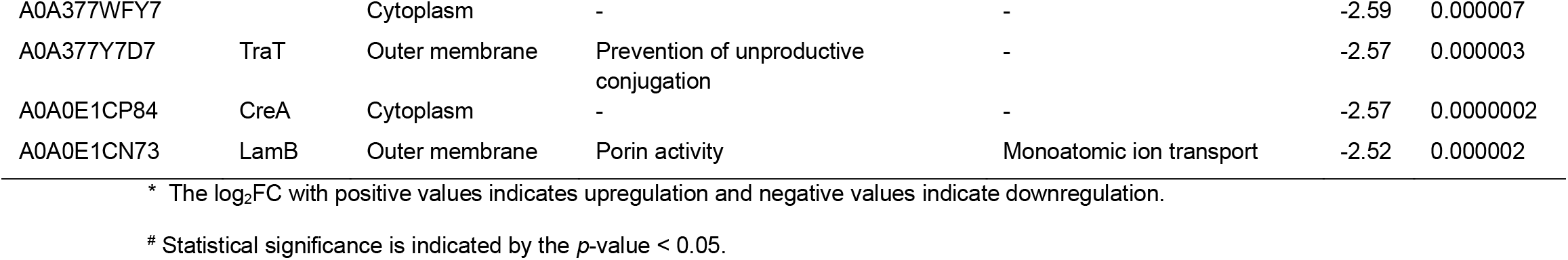
List of the top upregulated and downregulated proteins in *K. pneumoniae* ATCC 13883 treated with colistin.

**Fig. 2.**
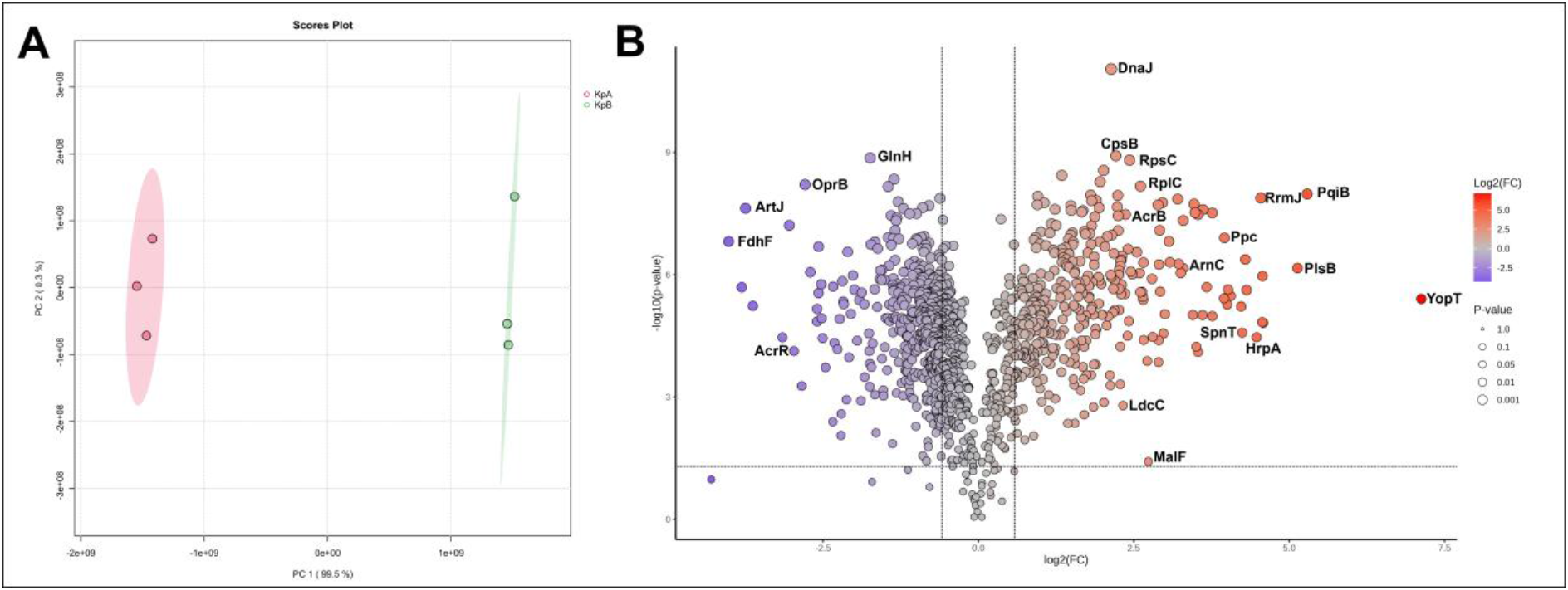
Quantitative proteomic profiling of *K. pneumoniae* in response to colistin exposure. (A) Principal Component Analysis (PCA) score plot showing the global proteomic perturbation between untreated control (KpA) and colistin (2 μg/ml) treated samples (KpB). The tight clustering within the 95% confidence intervals indicates high reproducibility and distinct protein expression profiles between the two groups. (B) Volcano plot of differentially abundant proteins (DAPs). The distribution of all identified proteins based on their fold change (log_2_FC) and statistical significance -log_10_ *p*-value ≤ 0.05. Red dots represent proteins significantly upregulated (FC ≥ 1.5), while blue dots represent proteins significantly downregulated (FC ≤ 0.66). Grey dots represent proteins that did not meet the significance threshold.

**Fig. 3.**
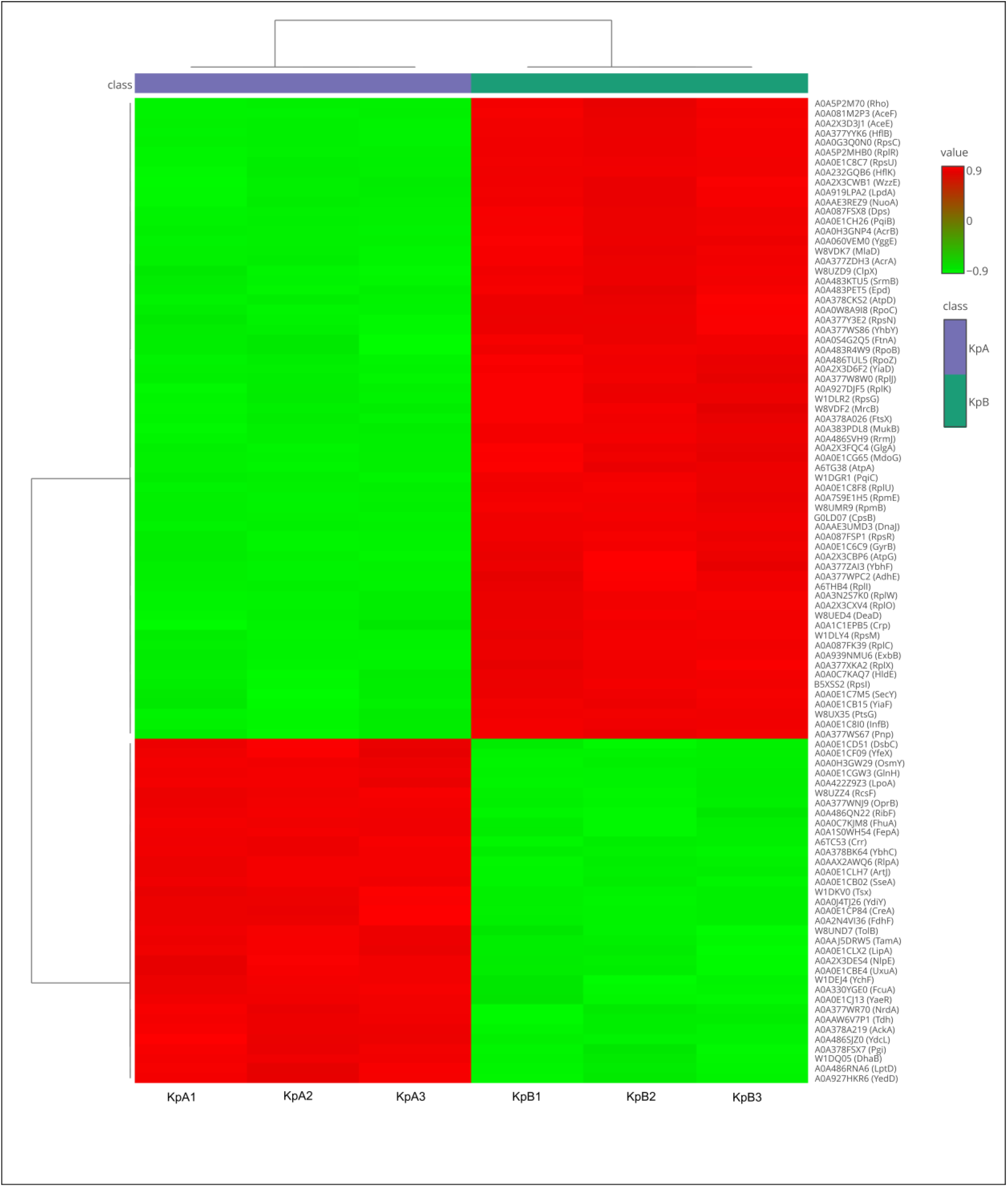
Expression profile of the top 100 DAPs. Hierarchical clustering and heatmap analysis of the 100 most significant DAPs (*p* ≤ 0.05) identified in *K. pneumoniae* in response to colistin exposure. Rows represent individual proteins, and columns represent biological replicates for the control (KpA) and colistin-treated (KpB) groups. The colour scale indicates relative protein abundance, with red indicating upregulation (FC ≥ 1.5) and green indicating downregulation (FC ≤ 0.66). The dendrogram on the left illustrates the degree of similarity in expression patterns among the selected proteins.

### Cellular localisation of the DAPs

Analysis of the 718 DAPs in colistin-treated *K. pneumoniae* revealed distinct patterns of cellular localisation (Fig. 4A), providing insights into the bacterial response mechanisms. The most highly represented cellular components were the cytosol, plasma membrane, periplasm, and outer membrane, followed by other cell components. The largest proportion of DAPs (n = 440, > 60%) was from the cytoplasm. The next most prevalent proteins (n = 173) based on localisation were bacterial membrane systems. Specifically, 130 proteins detected were from the plasma membrane. This strongly correlates with colistin’s mechanism of action. The localisation information for 57 proteins could not be determined, indicating a need for further characterisation of their precise subcellular localisation.

**Fig. 4.**
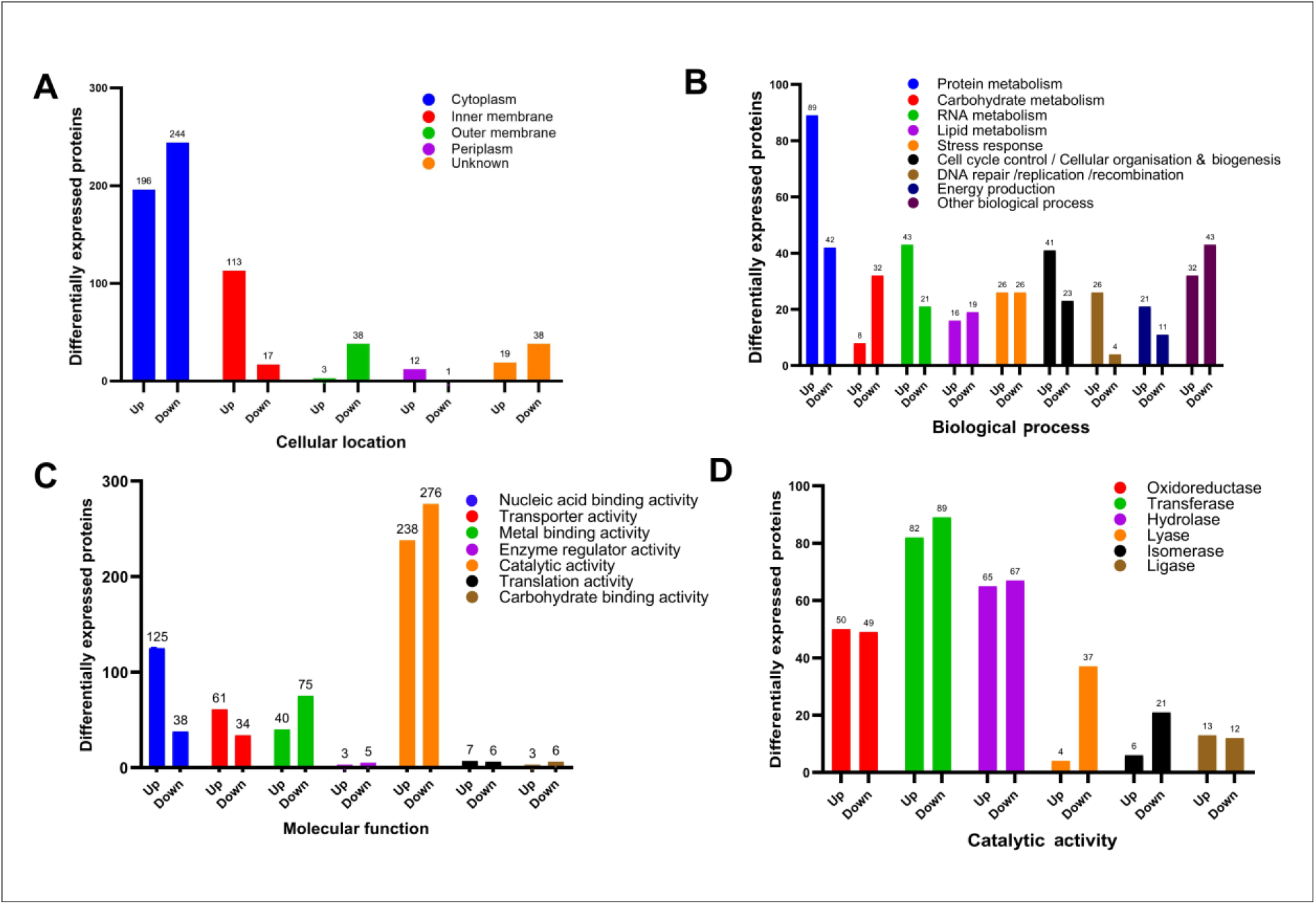
Gene ontology (GO) analysis of the DAPs. (A) Subcellular localisation. Bar charts representing the distribution of upregulated (n = 339) and downregulated (n = 379) proteins across different cellular compartments. (B) Biological Process. Comparative analysis of DAPs involved in key metabolic and cellular pathways. (C) Molecular Function. Classification of DAPs based on their molecular roles. (D) Catalytic activity. Breakdown of the various enzyme classes affected by colistin treatment.

Among the total of 339 upregulated proteins, 196 proteins were of cytoplasmic origin, whereas 113 were from the plasma membrane (Fig. 4A). Moreover, 12 upregulated proteins were of periplasmic origin, and 3 were from the outer membrane (Fig. 4A). Conversely, the analysis of 379 down-regulated proteins showed a few interesting insights. As with the upregulated set, cytoplasmic proteins (n = 244) were the most prominent among the downregulated ones. A notable contrast to the upregulated set was observed in membrane systems. The outer membrane showed a highly significant downregulation, with 38 proteins accounting for 9% of all down-regulated proteins (Fig. 4A). The plasma membrane showed a significantly lower number of down-regulated proteins (n = 17) than the upregulated ones. Similarly, minor compartments such as the fimbria, microcompartment, and extracellular spaces exhibited exclusive downregulation, indicating a shutdown of non-essential functions to conserve energy and resources for survival.

### Biological processes of the DAPs

The functional annotation of the DAPs revealed a robust, strategic reorganisation of the *K. pneumoniae* proteome in response to the colistin challenge, prioritising immediate survival mechanisms over cellular growth. The most significant increase was observed in proteins involved in protein and amino acid metabolism, showing a strong bias toward upregulation (89 upregulated vs. 42 downregulated). This may represent a stress response mechanism that repairs damage or produces new proteins necessary for survival. The transport category, which is key to maintaining homeostasis and directly interacting with the antibiotic, was among the most active processes, with 108 DAPs (58 up-regulated and 50 down-regulated). RNA metabolism and transcription were strongly upregulated (43 out of 64 DAPs), indicating increased transcriptional activity for adaptive gene expression (Fig. 4B). Additionally, DNA replication, repair, and recombination were predominantly activated (26 upregulated out of 30 DAPs), reflecting a focus on repairing genetic damage and preserving genome integrity under antibiotic stress (Fig. 4B). Proteins involved in energy production (32 DAPs) and cell cycle control and biogenesis (64 DAPs) were also significantly upregulated, suggesting that the cell was actively repairing damage, adapting to the hostile environment, and sustaining essential functions (Fig. 4B).

The stress response pathway, representing the cell’s direct mechanisms for sensing and mitigating colistin damage, contained 52 total DAPs, with a highly balanced expression profile: 26 proteins up-regulated and 26 down-regulated (Fig. 4B). Conversely, carbohydrate metabolism (40 DAPs) was the most strongly down-regulated (32 downregulated vs. 8 upregulated), indicating a shift away from standard growth-associated metabolic schemes to conserve energy (Fig. 4B).

### Molecular function of the DAPs

The molecular function classification of the DAPs demonstrates a clear strategic shift in the cellular machinery of *K. pneumoniae* to counter colistin. The most dominant functional group is catalytic activity, encompassing 514 total DAPs, a majority of which were downregulated (n = 276), indicating broad-scale suppression of several enzymatic reactions (Fig. 4C). In contrast, of the total 163 DAPs with nucleic acid binding activity detected, a large proportion (n = 125) were upregulated. Following this, of the 95 DAPs with transporter activity, 61 were found to be upregulated, indicating the functional basis for active efflux and the critical re-routing of molecular traffic across the membranes. Further evidence of an active response to colistin treatment was observed with the upregulation of 21 proteins associated with ribosome structure and ribosome biogenesis, ensuring rapid capacity to synthesise the necessary stress-response proteins (Suppl. Data Sheet 1). Finally, metal-binding activity (115 total DAPs) showed an overall downregulation (n = 75), outside of catalytic activity, which possibly relates to the general metabolic slowdown or to the strategic modification of cell envelope properties (Fig. 4C).

The functional analysis of DAPs related to catalytic activity revealed a significant distribution across all major enzyme classes. Transferase activity was the largest subset, with 171 DAPs: 89 downregulated and 82 upregulated. Further analysis among the transferases indicated that 31 DAPs have kinase activity, of which 23 were downregulated, and 8 were upregulated. Of the 132 DAPs with hydrolase activity, 67 were downregulated, and 65 were upregulated in response to colistin exposure. Other significant catalytic proteins are presented in Fig. 4D.

### Gene ontology (GO) enrichment analysis

#### Cellular component

The gene ontology (GO) term enrichment analysis for the cellular component category revealed significant over-representation of terms related to the ribosome and general organelle structure, as well as notable enrichment for membrane components. The highest enrichment and statistical significance were observed across five related terms: ribosome, organelle, intracellular organelle, non-membrane-bounded organelle, and intracellular non-membrane-bounded organelle (Fig. 5A). These terms were all associated with the largest group of genes, approximately 160. This dominant enrichment strongly suggests that the cellular location most affected by colistin exposure is the cell’s protein synthesis machinery. Furthermore, terms related to the cell boundary were also enriched, including the intrinsic component of the membrane (Fig. 5A). Therefore, these findings indicate that while ribosomal components are the primary cellular structures affected, changes to the cell membrane and its components also represent a significant cellular response.

**Figure 5.**
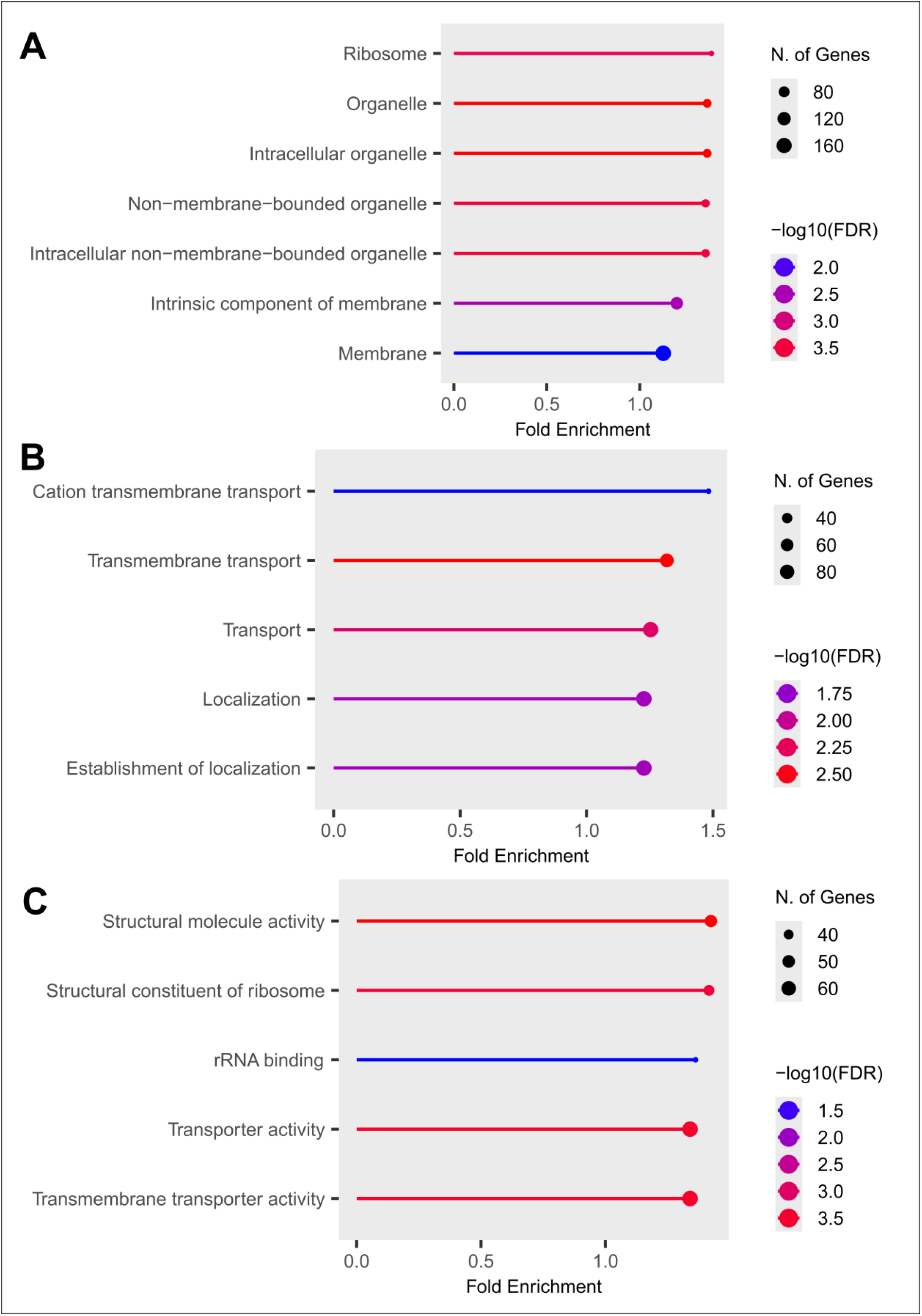
Gene Ontology (GO) enrichment analysis of the DAPs. The bar charts show statistically significant enrichment of GO terms across three categories. The X-axis represents fold enrichment, with statistical significance indicated by the colour gradient or –log_10_(FDR) values. (A) Cellular component. (B) Biological process. (C) Molecular function.

#### Biological processes

The GO enrichment analysis revealed that several key biological processes were significantly enriched, with a strong focus on cellular transport and localisation mechanisms. The most significantly enriched GO term was cation transmembrane transport, with a fold enrichment of approximately 1.5, a -log_10_(FDR) value of approximately 2.50, and involvement of over 20 genes (Fig. 5B). Similarly, transmembrane transport showed a 1.4-fold enrichment and involved approximately 80 genes. Other significantly enriched terms included transport (1.3-fold enrichment), localisation (1.25-fold enrichment), and establishment of localisation (1.2-fold enrichment) (Fig. 5B). These findings indicated a statistically significant dysregulation in membrane-associated transport functions, particularly those related to cation movement, suggesting that processes involved in cellular transport and the establishment of molecular localization represent the primary biological functions affected by colistin treatment.

#### Molecular function

The GO enrichment analysis for the molecular function category identified several key activities that were significantly enriched, with a strong focus on structural roles and transmembrane movement. The two most enriched terms, both with the highest observed fold enrichment of approximately 1.43, were structural molecule activity and structural constituent of ribosome (Fig. 5C). Each term was associated with approximately 60 genes. This indicates a significant impact on the components responsible for maintaining cellular architecture and the machinery of protein synthesis. Furthermore, the analysis highlighted roles in transport, specifically transmembrane transporter activity and the broader transporter activity, both exhibiting approximately 1.41-fold enrichment involving about 60 genes each. The term rRNA binding was also highly enriched (1.39-fold, 60 genes), further supporting the enrichment of ribosome-related functions (Fig. 5C). Collectively, these results suggest that the primary changes in the molecular functions of the system are driven by alterations in structural integrity, protein synthesis, and transport across cell membranes.

#### Significant DAPs

Proteomics analysis revealed that *K. pneumoniae* employs a survival strategy under colistin stress, characterised by significant upregulation of the arabinose operon proteins ArnC (log_2_FC = 3.29), ArnA (log_2_FC = 0.73), and ArnT (log_2_FC = 0.92), which are involved in the LPS modification pathway (Table 2). The highest upregulation was observed in YopT (log_2_FC = 7.12), followed by proteins involved in phospholipid and lipoprotein trafficking, such as PqiB (log_2_FC = 5.28) and PlsB (log_2_FC = 5.13) (Table 1).

**Table 2.** DAPs involved in various adaptive response pathways to colistin exposure identified in the study.

| Accession No.<br>(UniProt) | Protein | Description | log<sub>2</sub>FC | p-value | Functional outcome |
| --- | --- | --- | --- | --- | --- |
| LPS modification |  |  |  |  |  |
| A0A4P0YIA2 | ArnC | Undecaprenyl-phosphate 4-deoxy-4-formamido-L-arabinose transferase | 3.29 | 0.0000006 | Reduction of the overall negative charge in the outer membrane |
| A0A0E1CBB2 | ArnT | Undecaprenyl phosphate-alpha-4-amino-4-deoxy-L-arabinose arabinosyl transferase | 0.92 | 0.000007 |  |
| A0A378A1S4 | ArnA | Bifunctional polymyxin resistance protein | 0.73 | 0.0000007 |  |
| LPS biosynthesis |  |  |  |  |  |
| A0A3F3B9G4 | WaaA | 3-deoxy-D-manno-octulosonic acid transferase | 2.11 | 0.0003 | Stabilisation of LPS structure |
| A0A2X3CWB1 | WzzE | ECA polysaccharide chain length modulation protein | 1.52 | 0.0000002 |  |
| LPS transport |  |  |  |  |  |
| A0A377WHT4 | MsbA | Transport ATP-binding protein | 1.93 | 0.000005 | Maintenance of the outer membrane barrier under colistin stress |
| A0A377TJF6 | LptF | LPS export system permease protein | 1.33 | 0.0002 |  |
| A0A2X3CC60 | LptC | Multifunctional fusion protein | 0.62 | 0.008 |  |
| A0A0A5RLB5 | LptB | LPS export system ATP-binding protein | -0.77 | 0.000009 |  |
| A0A486RNA6 | LptD | LPS-assembly protein | -1.06 | 0.0000001 |  |
| Capsular polysaccharide synthesis |  |  |  |  |  |
| A0A1C3T1V8 | Wzc | Tyrosine-protein kinase | 4.00 | 0.000005 | Steric hindrance preventing colistin access to lipid A |
| Biofilm formation |  |  |  |  |  |
| A0A1Y0PX35 | RcsB | Transcriptional regulatory protein | 0.63 | 0.002 | Regulatory activation of protective envelope remodelling and biofilm-associated tolerance. |
| W8UDQ3 | OmpR | DNA-binding dual transcriptional regulator | 0.59 | 0.00004 |  |
| Efflux pump |  |  |  |  |  |
| A0A377ZDH3 | AcrA | RND family efflux transporter MFP subunit | 2.36 | 0.00000003 | Increased efflux activity |
| A0A0H3GNP4 | AcrB | Efflux pump membrane transporter | 2.88 | 0.00000001 |  |
| A0A377XIC4 | AcrR | Multidrug efflux pump AcrAB operon transcription repressor | -2.96 | 0.00007 |  |
| Porin channel |  |  |  |  |  |
| A0A0B7GG83 | OmpA | Outer membrane protein A | -0.80 | 0.000009 | Decreased outer membrane permeability |
| W8UKR5 | OmpX | Outer membrane protein X | -0.91 | 0.000001 |  |
| A0A0E1CN73 | LamB | Maltoporin | -2.52 | 0.0000017 |  |

The most prominent response to CAMP involved the synthesis of the capsular polysaccharide (CPS) and modification of the LPS (Table 2). Wzc, the tyrosine-protein kinase essential for capsule assembly and translocation, exhibited the highest degree of upregulation (log_2_FC = 4.00), accompanied by a significant increase in LPS-related enzymes, including WaaA (log_2_FC = 2.11) and WzzE (log_2_FC = 1.52) (Table 2).

The RND-family efflux pump components AcrB and AcrA were significantly upregulated by 2.88-fold and 2.36-fold, respectively (Table 2). This induction was strongly correlated with the suppression of the repressor AcrR (log_2_FC = -2.96) (Table 2). This was paradoxically accompanied by significant downregulation of the global activator RamA (log_2_FC = -0.76) and the outer membrane porin TolC (log_2_FC = -0.63). To support structural changes under stress, the Tol-Pal system, critical for maintaining OM stability, showed increased expression of TolQ (log_2_FC = 1.72) and TolA (log_2_FC = 1.43) upon RcsB induction (log_2_FC = 0.63) (Table 2).

To minimise polymyxin entry, *K. pneumoniae* significantly downregulated several major outer membrane proteins (OMPs). Key proteins suppressed included OmpA (log_2_FC = -0.80), OmpX (log_2_FC = -0.91), and the maltoporin LamB (log_2_FC = -2.52) (Table 2). Conversely, components of the LPS transport (Lpt) pathway showed divergent expression; while MsbA (log_2_FC = 1.93), LptF (log_2_FC = 1.33), and LptC (log_2_FC = 0.62) increased, LptB and LptD were downregulated to -0.77-fold and -1.06-fold, respectively, suggesting a potential regulatory bottleneck in membrane assembly (Table 2).

### Pathway enrichment analysis

The KEGG pathway enrichment analysis of *K. pneumoniae* under colistin stress revealed a synchronised metabolic shift characterised by the simultaneous downregulation of early-stage lipid A biosynthesis (LpxA, LpxC, LpxD) and the significant upregulation of the L-Ara4N modification pathway (ArnT) and core-attachment enzymes (WaaA), neutralising the negative charge of the OM to repel the cationic antibiotic (Fig. 6A; Table 2). This protective remodelling is fuelled by an intensified flux through the TCA cycle (Fig. S3), specifically via upregulated citrate synthase and isocitrate dehydrogenase, to meet the increased demand for ATP and biosynthetic precursors. Furthermore, the upregulation of peptidoglycan synthesis (MurA and PBPs) (Fig. S4) and biofilm-associated regulators (OmpR and RcsB) (Fig. S5) indicates an effort towards reinforcing cell wall integrity and transition into a protective biofilm state, while a strategic downregulation of ABC transporters and porins limits further antibiotic entry. The pathway analysis further reveals that *K. pneumoniae* can mount a robust defence against colistin through the Arn-mediated modification of Lipid A, even when the classical PhoPQ and PmrAB regulators are not the primary drivers (Fig. 6B).

**Figure 6.**
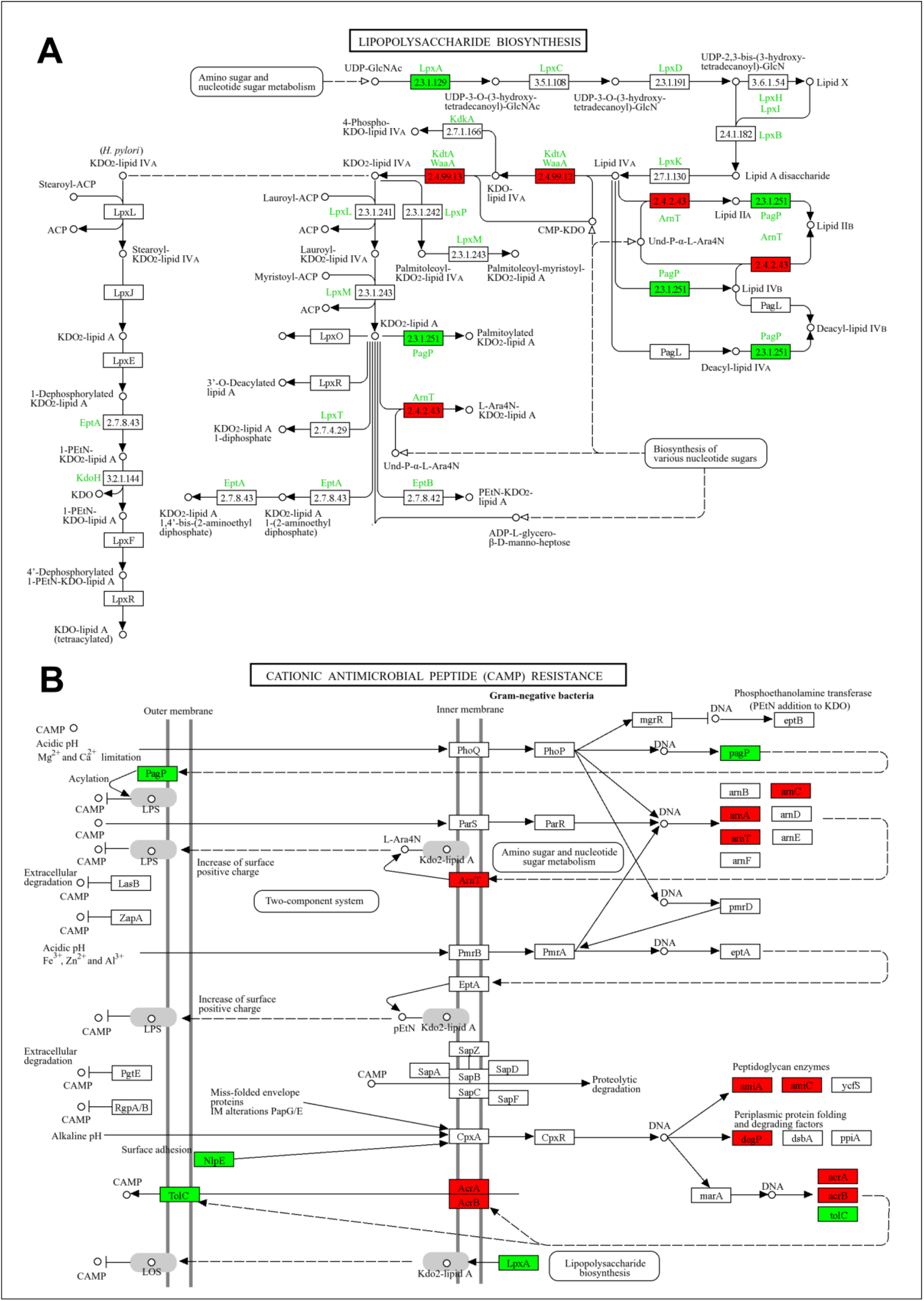
KEGG pathway enrichment analysis of the DPAs in response to colistin treatment. Red boxes/arrows indicate significantly upregulated pathways/proteins, while green indicates downregulation. (A) Lipopolysaccharide biosynthesis pathway. (B) Cationic Antimicrobial Peptide (CAMP) resistance pathway.

## Discussion

The significant rise in nosocomial infections caused by *K. pneumoniae*, especially following the COVID-19 pandemic, poses a major challenge in the healthcare sector. Moreover, the rising incidence of resistance to the frontline antibiotics, including carbapenems and polymyxins, among the opportunistic pathogens is a grave concern requiring urgent intervention. The molecular mechanisms underlying polymyxin resistance in *K. pneumoniae* are well understood. However, our knowledge of the adaptive response to polymyxin treatment in sensitive pathogens remains inadequate. This study aims to provide a comprehensive insight into the adaptive landscape of *K. pneumoniae* in response to polymyxin E (colistin), unveiling a complex survival-over-growth strategy coordinated through membrane remodelling and translational reprogramming.

In this study, we observed that *K. pneumoniae* cultures at mid-log phase (OD_600_ = 0.4) challenged with 2 µg/ml colistin initially reduced viable cells, followed by a significant increase in their number (Fig. 1C). This implies that significant number of susceptible cells are killed initially and in a short time the bacteria started challenging the antibiotic stress by activating the stress response pathways. The primary objective of this investigation was to understand the initial cellular response at the proteomics level. It is quite possible that the surviving bacteria have altered their cell membranes to reduce colistin binding or have activated the efflux pumps to expel the antibiotic from the cell [6,31].

As colistin targets the bacterial cell envelope, the shift in membrane protein abundance was the most notable response, with 113 plasma membrane proteins significantly upregulated, in contrast to the downregulation of 38 outer membrane proteins (Fig. 4A). The upregulation of transporter activity (95 DAPs) and cation transmembrane transport indicated the activation of the drug efflux or rebalancing the ionic gradient disturbed by polymyxin binding (Fig. 4C). The enrichment of cation transmembrane transport is crucial; by modulating the charge of the cell envelope through the addition of L-Ara4N or pEtN to lipid A, *K. pneumoniae* reduces its electrostatic affinity for the positively charged colistin molecules [10].

Previous studies have shown that, in colistin-resistant *K. pneumoniae, arnBCADTEF* operon expression is increased compared to the colistin-sensitive *K. pneumoniae*, through the activation of PhoPQ and PmrAB TCS [13,19]. Moreover, transcriptomic analysis reveals that activating mutations in the *crrB* sensor kinase lead to increased expression of the *arnBCADTEF* operon, a common feature identified in colistin-resistant clinical strains [32]. Interestingly, in our study, the expression of proteins encoded by the *arnBCADTEF* operon, including ArnC, ArnA, and ArnT, was significantly upregulated compared to the no-antibiotic-treated culture (Table 2). ArnT is a transferase enzyme that adds an L-Ara-4N molecule to the lipid A portion of the LPS. This addition provides resistance to colistin [33]. However, a notable change from previous studies was the detection of no significant changes in the expression of PhoPQ and PmrAB signalling proteins, suggesting that other proteins or TCS modules may be involved in the response mechanism. Fleitas et al. [14] have shown that ArnB and ArnC are upregulated in *K. pneumoniae* in response to the antimicrobial peptide PaDBS1R1, indicating their role in the general response to cell envelope stress. Interestingly, ArnT expression was reduced in the *K. pneumoniae* outer membrane vesicles (OMVs) proteome upon polymyxin exposure [16]. It was suggested that this could be due to the employment of OMVs with unmodified LPS as decoys for polymyxin binding, thereby protecting the bacterial cell with modified LPS from polymyxin [16,34].

Proteomics results indicated targeted remodelling of the LPS transport machinery with the identification of five key components of the Lpt machinery, LptB, LptD, LptF, LptC, and MsbA (Table 2) [35,36]. The upregulation of the inner membrane and periplasmic components MsbA, LptF, and LptC indicates the readiness of the system to export LPS components to repair membrane damage due to colistin exposure (Fig. 6A; Table 2). Moreover, we observed marked downregulation of the ATPase LptB and the OM translocon LptD of the Lpt system (Table 2). This pattern of proteomic perturbation suggests that the cell, while preparing to address membrane damage by stockpiling LPS transport intermediates at the inner membrane and restricting OM delivery, maintains a state of internal readiness for rapid envelope repair once the antimicrobial stress subsides.

Colistin treatment in *K. pneumoniae* revealed a distinct regulatory divergence in the AcrAB-TolC efflux system. While downregulation of the repressor AcrR led to the expected upregulation of the plasma membrane components AcrA and AcrB (Table 2), this paradoxically coincided with significant downregulation of the OM porin TolC (log_2_FC = -0.63) and the global activator RamA (log_2_FC = -0.76). Previous proteomics studies have also detected the upregulation of AcrA and AcrB proteins in response to sub-inhibitory concentrations of polymyxin B [16,33]. Consistent with our findings, the upregulation of AcrA and AcrB in colistin-resistant *K. pneumoniae* suggests that efflux-mediated mechanisms contribute to resistance in the absence of specific TCS mutations [37]. In our study, AcrR protein expression was downregulated under colistin treatment, a response consistent with other environmental stressors and antibiotics that trigger AcrR suppression to facilitate the efficient efflux of toxic substances [2]. The proteomics analysis revealed a significant upregulation of an ABC transporter and MDR efflux pump YbhF (log_2_FC = 2.03), which is often upregulated under envelope stress [38].

The study revealed that while the OM protein TolC was downregulated (log_2_FC = -0.63), there was significant upregulation of TolQ (log_2_FC = 1.72) and TolA (log_2_FC = 1.43) proteins (Suppl. data sheet 1). Colistin acts by displacing Mg^2+^ and Ca^2+^ ions from the LPS, causing the OM to bubble and pull away from the IM [39]. TolQ and TolA proteins are core components of the Tol-Pal system, which acts as a molecular staple that anchors the OM to the IM and the peptidoglycan layer [40]. When colistin damages the membrane, the Rcs stress response system is triggered. Rcs is known to upregulate TolQ and TolA to prevent membrane blebbing and stabilise the cell envelope [41]. Interestingly, our data revealed a downregulation of the global activator RamA, which is responsible for the observed decrease in TolC expression, as RamA is a known positive regulator of TolC [42]. TolC, a large, relatively open porin-like channel spanning the IM and OM, is energetically expensive [43]. In a high-stress environment, the cell may prioritise sealing the membrane over efflux [44]. By downregulating TolC, the bacteria reduce the number of porin channels through which the antibiotic could penetrate. Therefore, the observed downregulation of TolC alongside the upregulation of TolA and TolQ (envelope stabilisers) suggests a transition from an energy-intensive efflux-based defence to a structural sealing strategy [44].

Our study further revealed coordinated activation of the Rcs phosphorelay, characterised by upregulation of the response regulator RcsB (Table 2). This is functionally consistent with our observed upregulation of the TolA (log_2_FC = 1.43) and TolQ (log_2_FC = 1.72) proteins of the TolQRA operon, as RcsB is a known activator of the TolQRA operon in *E. coli* [45]. Interestingly, the observed downregulation of the sensor RcsF (log_2_FC = -1.32) suggests a potential feedback mechanism or a strategy to minimise the synthesis of non-essential OM lipoproteins during colistin-induced membrane destabilisation. Moreover, over-activation can lead to excessive capsule production, which could halt growth [46].

The proteomics investigation revealed a significant upregulation of the tyrosine kinase Wzc, which is involved in the synthesis of the capsule responsible for bacterial virulence and associated with the development of colistin resistance [47]. Interestingly, a previous phosphoproteomics study had identified Wzc as one of the three tyrosine kinases in *K. pneumoniae* [48]. The polyphosphate kinase Ppk is a central enzyme in the metabolism of inorganic polyphosphate (polyP), required for stress survival, formation of biofilm, capsule synthesis, and virulence [49,50]. Our proteomics analysis showed a significant upregulation of Ppk (log_2_FC = 1.89), possibly a response to the colistin treatment. Rojas et al. [50] investigated proteomic perturbation in a *K. pneumoniae ppk1* mutant background and found the differential expression of proteins involved in capsule synthesis, biofilm formation, and lipid A modification.

In Gram-negative bacteria, the integrity of the cell envelope is maintained by a sophisticated network of signalling pathways, including the five most extensively characterised envelope stress response (ESR) systems, namely the Psp (phage shock protein), Cpx (conjugative pilus expression), σ^E^ (sigma E), Bae (bacterial adaptive response), and Rcs (regulator of capsule synthesis) pathways. These systems are defined by the coordinated upregulation of specific proteomes and chaperones to mitigate damage to the inner membrane, periplasm, or outer membrane [31]. In the current study, colistin treatment significantly modulated the bacterial proteome associated with the ESR system, characterised by marked upregulation of PspA (log_2_FC = 2.59) and DegS (log_2_FC = 2.26) (Suppl. Data Sheet 1).

One of the most striking findings was the upregulation of all 21 proteins associated with ribosome biogenesis, along with a high enrichment of ribosomal structural constituents (Suppl. Data Sheet 1). This indicates the rapid synthesis of stress-response proteins, such as chaperones and repair enzymes, identified in our biological function analysis. To fuel this energy-intensive repair process, the bacteria appear to undergo a metabolic shift. The strong down-regulation of carbohydrate metabolism (32 DAPs) and catalytic activity (276 DAPs) suggests a strategic quiescence (Figs. 4B, 4C, 4D). By suppressing standard growth-related metabolic pathways, the cell conserves resources for essential survival functions, such as DNA repair and membrane maintenance.

The molecular function analysis of upregulated proteins reveals that transporter activity is a key feature across multiple cellular compartments, particularly pronounced within membrane systems. This strong emphasis on membrane-associated transporters underscores their vital role in potentially facilitating the efflux of colistin. Additionally, the significant presence of kinase activity among membrane proteins suggests the activation of sensory and signalling networks at the cell surface, which are crucial for orchestrating the bacterial response to antibiotic stress (Fig. 4D).

## Conclusion

This study suggests that *K. pneumoniae* responds to colistin exposure through a highly coordinated reorganisation of cellular resources to ensure its survival. This is primarily achieved by outer membrane remodelling, through the significant upregulation of the *arnBCADTEF* operon. The findings further indicate a PhoPQ/PmrAB-independent activation of the LPS modification pathways, which warrants further investigation. Moreover, the identification of DAPs involved in CAMP resistance, biofilm formation, capsular synthesis, and peptidoglycan synthesis pathways suggests the multifrontal response mounted by the bacteria against colistin exposure. In conclusion, this work provides insights into a host of regulatory pathways perturbed by colistin treatment, which could be exploited to identify novel drug targets and explore synergistic drug combinations to dismantle bacterial response mechanisms and restore colistin’s effectiveness.

## Supporting information

Supplementary Data Sheet

## Acknowledgements

We express our sincere gratitude to Dr. Harapriya Mohapatra, NISER, Bhubaneswar, for the *K. pneumoniae* strain; to Ms. Harsha Sankar, IISER, Berhampur, for her technical support with ultracentrifugation; and to Mr. R. Rajendra Reddy, Institute of Life Sciences, Bhubaneswar, for his help with proteomics work and data analysis. We acknowledge the infrastructural support from the Department of Biotechnology, Berhampur University.

## Author contributions

Conceptualisation: SKD and SSM; Data extraction and analysis: SKD, IP, and SSM; Data validation: SKD, IP, and SSM; Manuscript writing and reviewing: SKD, SKP, and SSM. All authors agreed to the submitted version.

## Data availability

The mass spectrometry proteomics data have been deposited in the ProteomeXchange Consortium (https://www.proteomexchange.org) via the PRIDE partner repository, with the dataset identifier PXD076091 and DOI 10.6019/PXD076091.

## References

[1] K.L. Wyres, M.M.C. Lam, K.E. Holt, Population genomics of Klebsiella pneumoniae, Nat. Rev. Microbiol. 18 (2020) 344–359. 10.1038/s41579-019-0315-1.

[2] L. Ding, S. Shen, J. Chen, Z. Tian, Q. Shi, R. Han, Y. Guo, F. Hu, Klebsiella pneumoniae carbapenemase variants: the new threat to global public health, Clin. Microbiol. Rev. 36 (2023) e00008–23. 10.1128/cmr.00008-23.

[3] W.R. Miller, C.A. Arias, ESKAPE pathogens: antimicrobial resistance, epidemiology, clinical impact and therapeutics, Nat. Rev. Microbiol. 22 (2024) 598–616. 10.1038/s41579-024-01054-w.

[4] S.S. Mohapatra, S.K. Dwibedy, I. Padhy, Polymyxins, the last-resort antibiotics: Mode of action, resistance emergence, and potential solutions, J. Biosci. 46 (2021) 85. 10.1007/s12038-021-00209-8.

[5] L. Poirel, A. Jayol, P. Nordmann, Polymyxins: Antibacterial Activity, Susceptibility Testing, and Resistance Mechanisms Encoded by Plasmids or Chromosomes, Clin. Microbiol. Rev. 30 (2017) 557–596. 10.1128/CMR.00064-16.

[6] I. Padhy, S.K. Dwibedy, S.S. Mohapatra, A molecular overview of the polymyxin-LPS interaction in the context of its mode of action and resistance development, Microbiol. Res. 283 (2024) 127679. 10.1016/j.micres.2024.127679.

[7] S.K. Dwibedy, I. Padhy, S.S. Mohapatra, A comprehensive analysis of the mutational landscape determining polymyxin resistance in Klebsiella pneumoniae, J. Antimicrob. Chemother. 81 (2026) dkaf503. 10.1093/jac/dkaf503.

[8] S.K. Dwibedy, I. Padhy, A.K. Panda, S.S. Mohapatra, Colistin resistance among the Gramnegative nosocomial pathogens in India: a systematic review and meta-analysis, J. Chemother. (2024) 1–13. 10.1080/1120009X.2024.2405355.

[9] E.V.K. Ledger, A. Sabnis, A.M. Edwards, Polymyxin and lipopeptide antibiotics: membrane-targeting drugs of last resort: This article is part of the Bacterial Cell Envelopes collection., Microbiology 168 (2022). 10.1099/mic.0.001136.

[10] L. Sun, Y. Zhang, T. Cai, X. Li, N. Li, Z. Xie, F. Yang, X. You, CrrAB regulates PagP-mediated glycerophosphoglycerol palmitoylation in the outer membrane of Klebsiella pneumoniae, J. Lipid Res. 63 (2022) 100251. 10.1016/j.jlr.2022.100251.

[11] A. Samantha, A. Vrielink, Lipid A Phosphoethanolamine Transferase: Regulation, Structure and Immune Response, J. Mol. Biol. 432 (2020) 5184–5196. 10.1016/j.jmb.2020.04.022.

[12] S.C. Nang, J. Li, T. Velkov, The rise and spread of mcr plasmid-mediated polymyxin resistance, Crit. Rev. Microbiol. 45 (2019) 131–161. 10.1080/1040841X.2018.1492902.

[13] X. Chen, J. Tian, C. Luo, X. Wang, X. Li, M. Wang, Cell Membrane Remodeling Mediates Polymyxin B Resistance in Klebsiella pneumoniae: An Integrated Proteomics and Metabolomics Study, Front. Microbiol. 13 (2022) 810403. 10.3389/fmicb.2022.810403.

[14] O. Fleitas, W. Fontes, C.M. De Souza, M.C. Da Costa, M.H. Cardoso, M.S. Castro, M.V. Sousa, C.A.O. Ricart, M.H.S. Ramada, H.M. Duque, W.F. Porto, O.N. Silva, O.L. Franco, A proteomic perspective on the resistance response of Klebsiella pneumoniae to antimicrobial peptide PaDBS1R1, J. Antimicrob. Chemother. 79 (2024) 112–122. 10.1093/jac/dkad354.

[15] M.C. Goodyear, L. Seidel, J.R. Krieger, J. Geddes-McAlister, R.C. Levesque, C.M. Khursigara, Quantitative proteomics reveals unique responses to antimicrobial treatments in clinical Pseudomonas aeruginosa isolates, mSystems 8 (2023) e00491–23. 10.1128/msystems.00491-23.

[16] M. Hussein, R. Jasim, H. Gocol, M. Baker, V.J. Thombare, J. Ziogas, A. Purohit, G.G. Rao, J. Li, T. Velkov, Comparative Proteomics of Outer Membrane Vesicles from Polymyxin-Susceptible and Extremely Drug-Resistant Klebsiella pneumoniae, mSphere 8 (2023) e00537–22. 10.1128/msphere.00537-22.

[17] A. Rocker, J.A. Lacey, M.J. Belousoff, J.J. Wilksch, R.A. Strugnell, M.R. Davies, T. Lithgow, Global Trends in Proteome Remodeling of the Outer Membrane Modulate Antimicrobial Permeability in Klebsiella pneumoniae, mBio 11 (2020) e00603–20. 10.1128/mBio.00603-20.

[18] R. Jasim, M.A. Baker, Y. Zhu, M. Han, E.K. Schneider-Futschik, M. Hussein, D. Hoyer, J. Li, T. Velkov, A Comparative Study of Outer Membrane Proteome between Paired Colistin-Susceptible and Extremely Colistin-Resistant Klebsiella pneumoniae Strains, ACS Infect. Dis. 4 (2018) 1692– 1704. 10.1021/acsinfecdis.8b00174.

[19] C.H.P. Cheung, P. Dulyayangkul, K.J. Heesom, M.B. Avison, Proteomic Investigation of the Signal Transduction Pathways Controlling Colistin Resistance in Klebsiella pneumoniae, Antimicrob. Agents Chemother. 64 (2020) e00790–20. 10.1128/AAC.00790-20.

[20] F. Fan, G. Chen, S. Deng, L. Wei, Proteomic analysis of meropenem-induced outer membrane vesicles released by carbapenem-resistant Klebsiella pneumoniae, Microbiol. Spectr. 12 (2024) e02917–23. 10.1128/spectrum.02917-23.

[21] S.K. Dwibedy, I. Padhy, G.M. Bitode, S.S. Mohapatra, Prevalence of polymyxin heteroresistance in Klebsiella pneumoniae under sustained antibiotic selection during experimental evolution, Arch. Microbiol. 207 (2025) 201. 10.1007/s00203-025-04405-0.

[22] CLSI, Performance Standards for Antimicrobial Susceptibility Testing, in: P.A. Wayne (Ed.), CLSI Suppl. M100, 30th ed., Clinical and Laboratory Standards Institute, 2020.

[23] N.J. Kruger, The Bradford Method For Protein Quantitation, in: Protein Protoc. Handb., Humana Press, Totowa, NJ, 2009: pp. 17–24. 10.1007/978-1-59745-198-7_4.

[24] V. Suresh, V. Mohanty, K. Avula, A. Ghosh, B. Singh, R.K. Reddy, D. Parida, A.R. Suryawanshi, S.K. Raghav, S. Chattopadhyay, P. Prasad, R.K. Swain, R. Dash, A. Parida, G.H. Syed, S. Senapati, Quantitative proteomics of hamster lung tissues infected with SARS-CoV-2 reveal host factors having implication in the disease pathogenesis and severity, FASEB J. 35 (2021) e21713. 10.1096/fj.202100431R.

[25] S. Behera, R.R. Reddy, K. Taunk, S. Rapole, R.R. Pharande, A.R. Suryawanshi, Delineation of altered brain proteins associated with furious rabies virus infection in dogs by quantitative proteomics, J. Proteomics 253 (2022) 104463. 10.1016/j.jprot.2021.104463.

[26] S. Jajula, V. Naik, B. Kalita, U. Yanamandra, S. Sharma, T. Chatterjee, S. Bhanuse, P.P. Bhavsar, K. Taunk, S. Rapole, Integrative proteome analysis of bone marrow interstitial fluid and serum reveals candidate signature for acute myeloid leukemia, J. Proteomics 303 (2024) 105224. 10.1016/j.jprot.2024.105224.

[27] Z. Pang, G. Zhou, J. Ewald, L. Chang, O. Hacariz, N. Basu, J. Xia, Using MetaboAnalyst 5.0 for LC–HRMS spectra processing, multi-omics integration and covariate adjustment of global metabolomics data, Nat. Protoc. 17 (2022) 1735–1761. 10.1038/s41596-022-00710-w.

[28] S.X. Ge, D. Jung, R. Yao, ShinyGO: a graphical gene-set enrichment tool for animals and plants, Bioinformatics 36 (2020) 2628–2629. 10.1093/bioinformatics/btz931.

[29] M. Kanehisa, M. Furumichi, Y. Sato, Y. Matsuura, M. Ishiguro-Watanabe, KEGG: biological systems database as a model of the real world, Nucleic Acids Res. 53 (2025) D672–D677. 10.1093/nar/gkae909.

[30] N.Y. Yu, J.R. Wagner, M.R. Laird, G. Melli, S. Rey, R. Lo, P. Dao, S.C. Sahinalp, M. Ester, L.J. Foster, F.S.L. Brinkman, PSORTb 3.0: improved protein subcellular localization prediction with refined localization subcategories and predictive capabilities for all prokaryotes, Bioinformatics 26 (2010) 1608–1615. 10.1093/bioinformatics/btq249.

[31] C. Brand, M. Newton-Foot, M. Grobbelaar, A. Whitelaw, Antibiotic-induced stress responses in Gram-negative bacteria and their role in antibiotic resistance, J. Antimicrob. Chemother. 80 (2025) 1165–1184. 10.1093/jac/dkaf068.

[32] M.S. Wright, Y. Suzuki, M.B. Jones, S.H. Marshall, S.D. Rudin, D. Van Duin, K. Kaye, M.R. Jacobs, R.A. Bonomo, M.D. Adams, Genomic and Transcriptomic Analyses of Colistin-Resistant Clinical Isolates of Klebsiella pneumoniae Reveal Multiple Pathways of Resistance, Antimicrob. Agents Chemother. 59 (2015) 536–543. 10.1128/AAC.04037-14.

[33] A.C.R. Lucena, M.G. Ferrarini, W.K. De Oliveira, B.H. Marcon, L.G. Morello, L.R. Alves, H. Faoro, Modulation of Klebsiella pneumoniae Outer Membrane Vesicle Protein Cargo under Antibiotic Treatment, Biomedicines 11 (2023) 1515. 10.3390/biomedicines11061515.

[34] J. Park, M. Kim, B. Shin, M. Kang, J. Yang, T.K. Lee, W. Park, A novel decoy strategy for polymyxin resistance in Acinetobacter baumannii, eLife 10 (2021) e66988. 10.7554/eLife.66988.

[35] P. Sperandeo, A.M. Martorana, A. Polissi, The lipopolysaccharide transport (Lpt) machinery: A nonconventional transporter for lipopolysaccharide assembly at the outer membrane of Gramnegative bacteria, J. Biol. Chem. 292 (2017) 17981–17990. 10.1074/jbc.R117.802512.

[36] F. Thélot, B.J. Orlando, Y. Li, M. Liao, High-resolution views of lipopolysaccharide translocation driven by ABC transporters MsbA and LptB2FGC, Curr. Opin. Struct. Biol. 63 (2020) 26–33. 10.1016/j.sbi.2020.03.005.

[37] S. Naha, K. Sands, S. Mukherjee, C. Roy, M.J. Rameez, B. Saha, S. Dutta, T.R. Walsh, S. Basu, KPC-2-producing Klebsiella pneumoniae ST147 in a neonatal unit: Clonal isolates with differences in colistin susceptibility attributed to AcrAB-TolC pump, Int. J. Antimicrob. Agents 55 (2020) 105903. 10.1016/j.ijantimicag.2020.105903.

[38] Z. Feng, D. Liu, L. Wang, Y. Wang, Z. Zang, Z. Liu, B. Song, L. Gu, Z. Fan, S. Yang, J. Chen, Y. Cui, A Putative Efflux Transporter of the ABC Family, YbhFSR, in Escherichia coli Functions in Tetracycline Efflux and Na+(Li+)/H+ Transport Front. Microbiol. 11 (2020) 556. 10.3389/fmicb.2020.00556.

[39] A. Sabnis, K.L. Hagart, A. Klöckner, M. Becce, L.E. Evans, R.C.D. Furniss, D.A. Mavridou, R. Murphy, M.M. Stevens, J.C. Davies, G.J. Larrouy-Maumus, T.B. Clarke, A.M. Edwards, Colistin kills bacteria by targeting lipopolysaccharide in the cytoplasmic membrane, eLife 10 (2021) e65836. 10.7554/eLife.65836.

[40] J. Szczepaniak, M.N. Webby, The Tol Pal system integrates maintenance of the three layered cell envelope, Npj Antimicrob. Resist. 2 (2024) 46. 10.1038/s44259-024-00065-0.

[41] J. Szczepaniak, C. Press, C. Kleanthous, The multifarious roles of Tol-Pal in Gram-negative bacteria, FEMS Microbiol. Rev. 44 (2020) 490–506. 10.1093/femsre/fuaa018.

[42] E.M. Grimsey, N. Weston, V. Ricci, J.W. Stone, L.J.V. Piddock, Overexpression of RamA, Which Regulates Production of the Multidrug Resistance Efflux Pump AcrAB-TolC, Increases Mutation Rate and Influences Drug Resistance Phenotype, Antimicrob. Agents Chemother. 64 (2020) e02460–19. 10.1128/AAC.02460-19.

[43] V. Koronakis, J. Eswaran, C. Hughes, Structure and Function of TolC: The Bacterial Exit Duct for Proteins and Drugs, Annu. Rev. Biochem. 73 (2004) 467–489. 10.1146/annurev.biochem.73.011303.074104.

[44] R.L. Guest, T.L. Raivio, Role of the Gram-Negative Envelope Stress Response in the Presence of Antimicrobial Agents, Trends Microbiol. 24 (2016) 377–390. 10.1016/j.tim.2016.03.001.

[45] Z. Li, Y. Zhu, W. Zhang, W. Mu, Rcs signal transduction system in Escherichia coli: Composition, related functions, regulatory mechanism, and applications, Microbiol. Res. 285 (2024) 127783. 10.1016/j.micres.2024.127783.

[46] L. Xu, J. Li, W. Wu, X. Wu, J. Ren, Klebsiella pneumoniae capsular polysaccharide: Mechanism in regulation of synthesis, virulence, and pathogenicity, Virulence 15 (2024) 2439509. 10.1080/21505594.2024.2439509.

[47] S. Khadka, B.E. Ring, R.S. Walker, L.R. Krzeminski, D.A. Pariseau, M. Hathaway, H.L.T. Mobley, L.A. Mike, Urine-mediated suppression of Klebsiella pneumoniae mucoidy is counteracted by spontaneous Wzc variants altering capsule chain length, mSphere 8 (2023) e0028823. 10.1128/msphere.00288-23.

[48] M.-H. Lin, T.-L. Hsu, S.-Y. Lin, Y.-J. Pan, J.-T. Jan, J.-T. Wang, K.-H. Khoo, S.-H. Wu, Phosphoproteomics of Klebsiella pneumoniae NTUH-K2044 Reveals a Tight Link between Tyrosine Phosphorylation and Virulence *, Mol. Cell. Proteomics 8 (2009) 2613–2623. 10.1074/mcp.M900276-MCP200.

[49] M.J. Gray, Inorganic Polyphosphate Accumulation in Escherichia coli Is Regulated by DksA but Not by (p)ppGpp, J. Bacteriol. 201 (2019) e00664–18. 10.1128/JB.00664-18.

[50] D. Rojas, A.E. Marcoleta, M. Gálvez-Silva, M.A. Varas, M. Díaz, M. Hernández, C. Vargas, G. Nourdin-Galindo, E. Koch, P. Saldivia, J. Vielma, Y.-H. Gan, Y. Chen, N. Guiliani, F.P. Chávez, Inorganic Polyphosphate Affects Biofilm Assembly, Capsule Formation, and Virulence of Hypervirulent ST23 Klebsiella pneumoniae, ACS Infect. Dis. 10 (2024) 606–623. 10.1021/acsinfecdis.3c00509.

