## Supplementary Figure for "Insights into the *Klebsiella pneumoniae* adaptive response mechanisms to colistin exposure using a label-free quantitative proteomics approach"

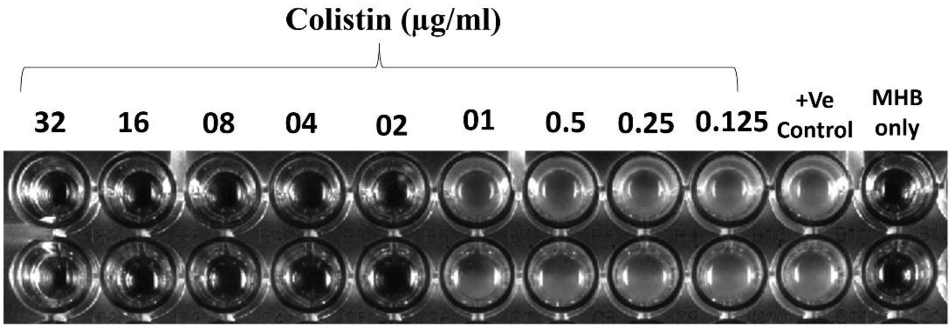


**Fig. S1.** Determination of the colistin MIC against *K. pneumoniae* ATCC 13883 by the broth microdilution (BMD) method. BMD assay showing the inhibitory effect of colistin at concentrations ranging from 0.125 µg/ml to 32 µg/ml. Bacterial growth is evident by turbidity at lower concentrations and in the positive control (+Ve), while the MIC is identified at 2 µg/ml, at which visible growth is first inhibited. MHB only serves as the sterility control.

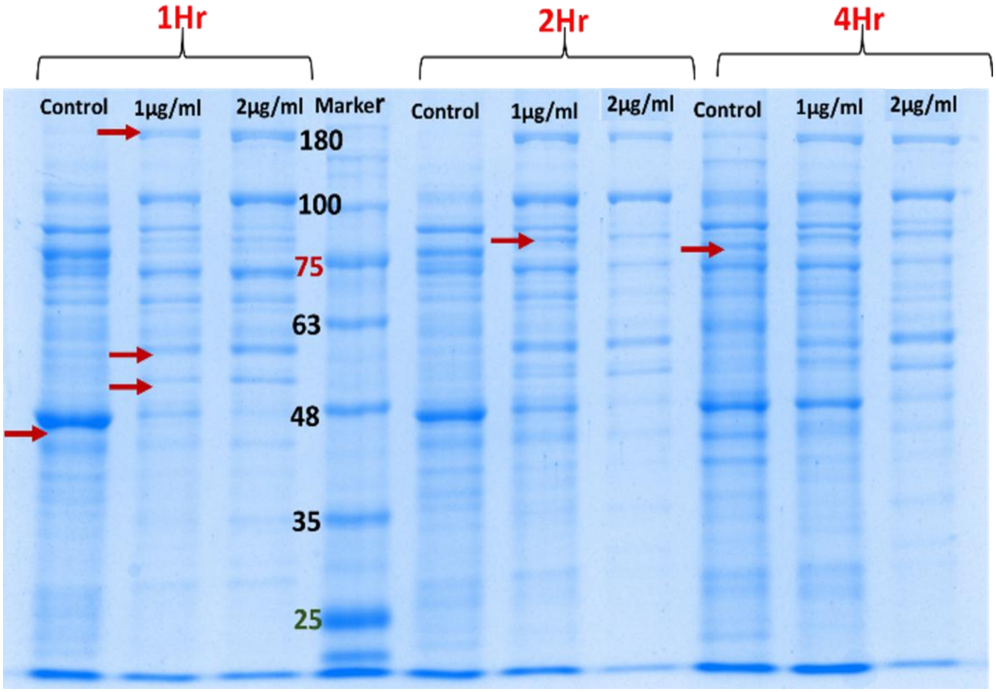


**Fig. S2.** SDS-PAGE profiling of membrane proteins prepared from the colistin-treated and untreated *K. pneumoniae* cultures. Extracted membrane proteins at 1, 2, and 4 hours post-exposure to colistin at sub-MIC (1µg/ml) and MIC (2 µg/ml) were performed on a 10% SDS-PAGE gel. Red arrows indicate a few differentially expressed protein bands in colistin-treated samples compared to the untreated controls.

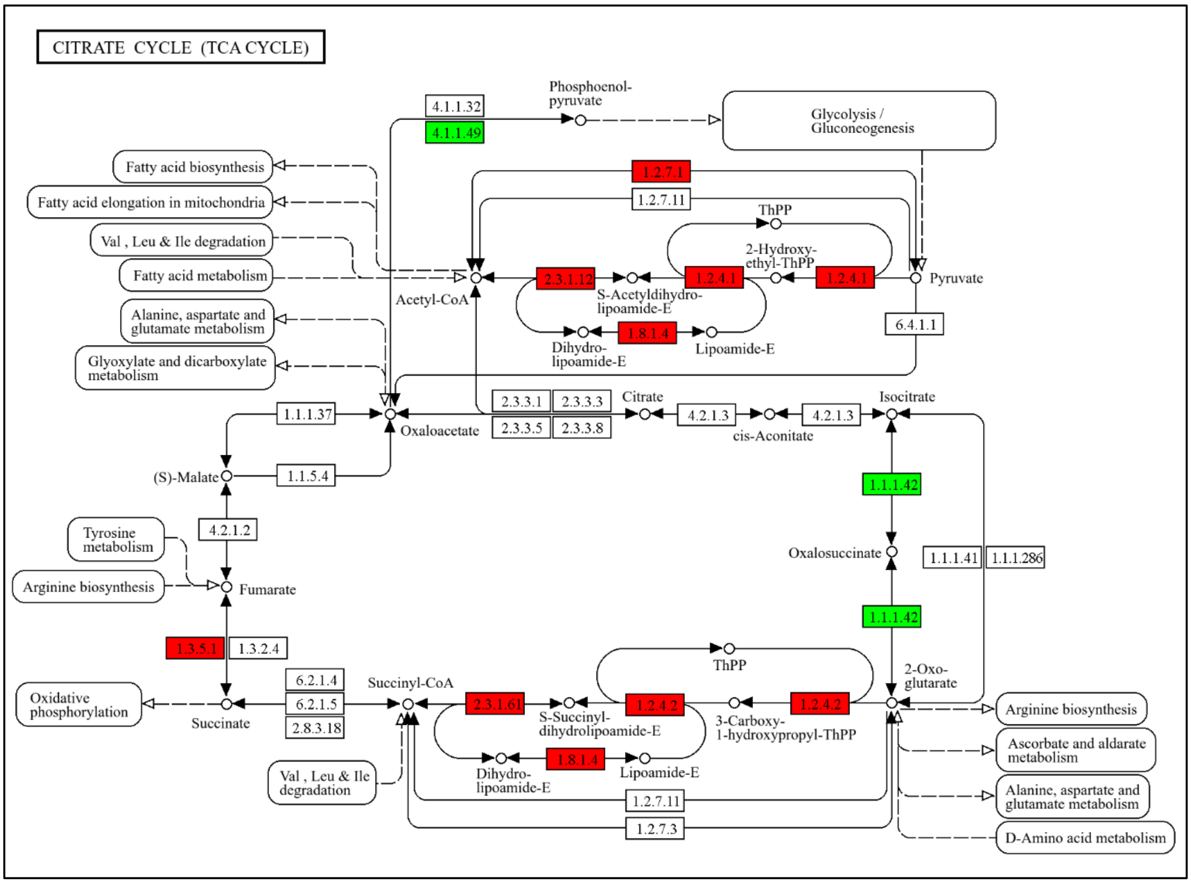


**Fig. S3.** TCA cycle flux during colistin-induced stress. KEGG pathway mapping of DAPs involved in the TCA cycle. Red boxes indicate significantly upregulated enzymes, including pyruvate dehydrogenase and succinate dehydrogenase, supporting an intensified metabolic flux to meet the high energy demands of early adaptive resistance mechanisms. Green boxes represent downregulated proteins.

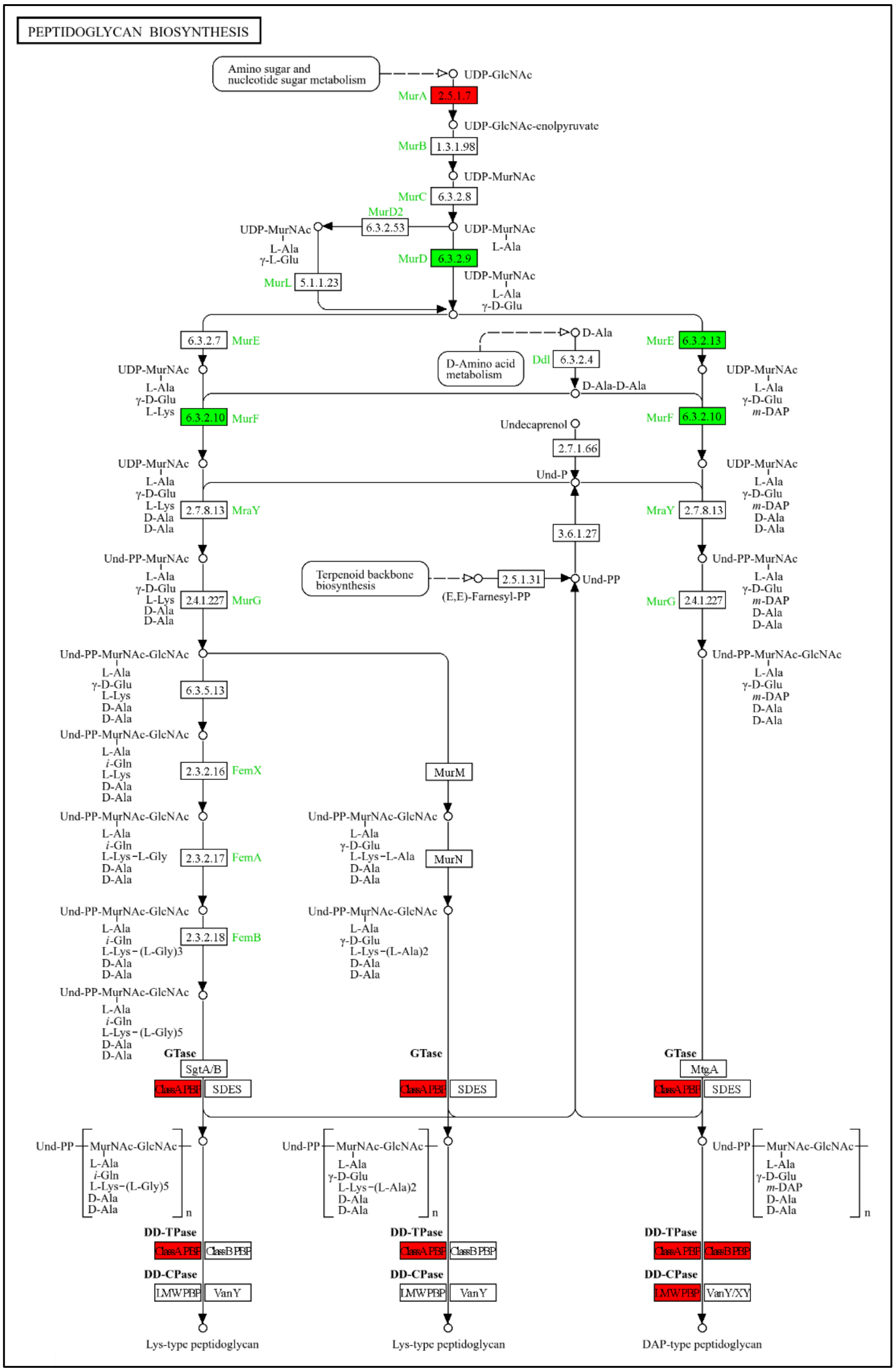


**Fig. S4.** Peptidoglycan biosynthesis pathway. KEGG pathway mapping of the DAPs involved in peptidoglycan synthesis and assembly. The coloured enzymes (red: upregulated; green: downregulated) show a complex restructuring of the cell wall. Significant upregulation of late-stage assembly proteins (e.g., penicillin-binding proteins/PBPs) suggests a structural reinforcement of the cell envelope to counter colistin-mediated membrane disruption.

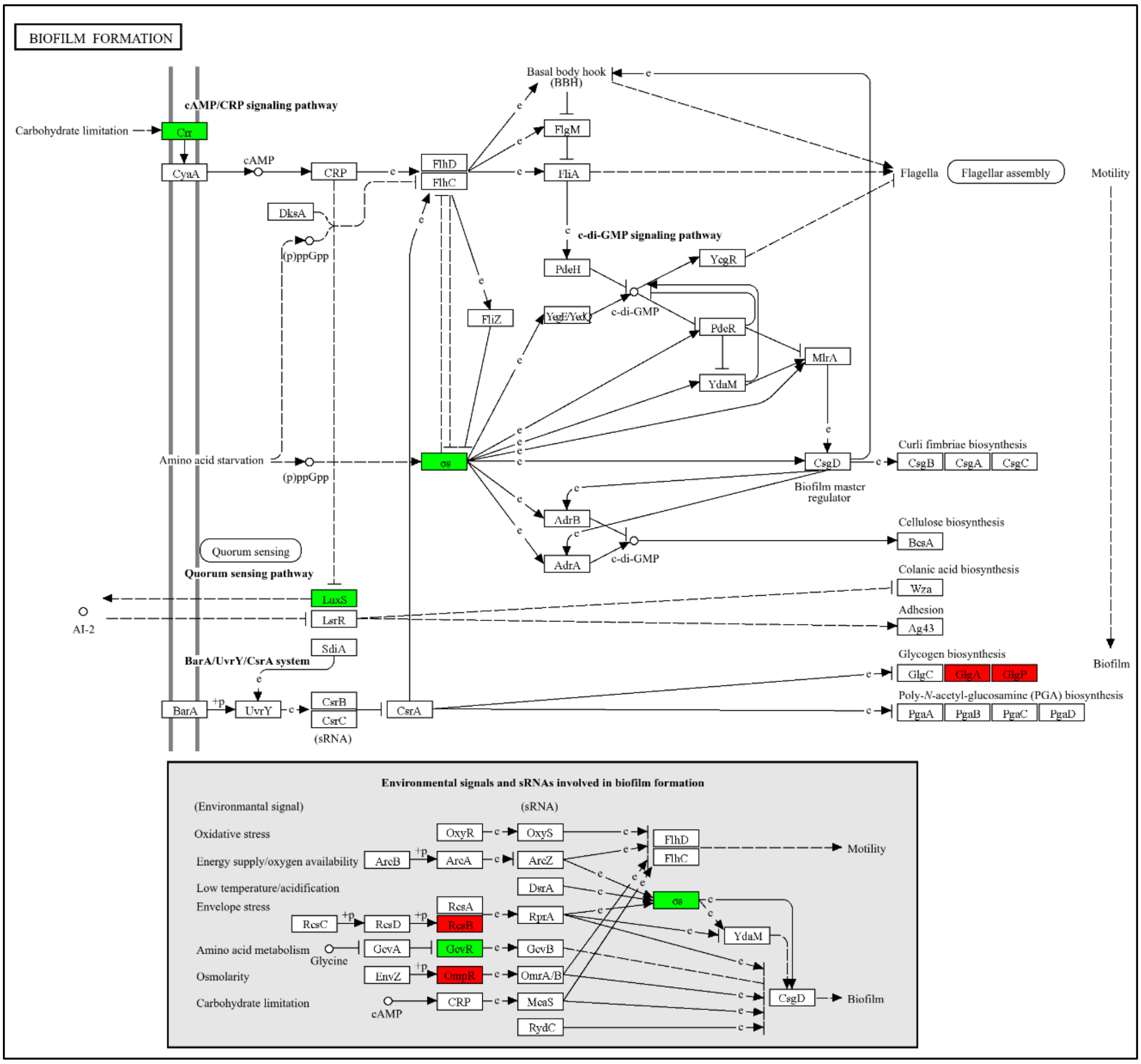


**Fig. S5:** Biofilm formation pathway. KEGG pathway mapping of the DAPs involved in biofilm regulation and signalling. Red indicates the upregulation of stress-responsive regulators (e.g., RcsB, OmpR), and green indicates downregulated proteins involved in the biofilm formation pathway.
